# Global proteomic profiling of primary macrophages during *M. tuberculosis* infection identifies TAX1BP1 as a mediator of autophagy targeting

**DOI:** 10.1101/534917

**Authors:** Jonathan M. Budzik, Nick E. Garelis, Teresa Repasy, Allison W. Roberts, Lauren M. Popov, Trevor J. Parry, David Jiminez-Morales, Danielle L. Swaney, Jeffrey R. Johnson, Nevan J. Krogan, Jeffery S. Cox

**Affiliations:** Department of Molecular and Cell Biology, University of California, Berkeley, Berkeley, CA, 94720; Department of Medicine, Division of Pulmonary and Critical Care Medicine, University of California, San Francisco, San Francisco, CA 94131; Department of Microbiology and Immunology, Program in Microbial Pathogenesis and Host Defense, University of California, San Francisco, San Francisco, CA 94158; Department of Medicine, Division of Cardiovascular Medicine, Stanford University, CA 94305, USA; Department of Cellular and Molecular Pharmacology, University of California, San Francisco, San Francisco, CA 94158

## Abstract

Macrophages are highly plastic cells that adopt diverse functional capabilities and play critical roles in immunity, cancer, and tissue homeostasis, but how these different cell fates and activities are triggered in response to their environmental cues is not well understood. We used new proteomic tools to identify protein post-translational modifications (PTMs) that control antibacterial responses of macrophages. Here, we report an unbiased and global analysis of the changes in host protein abundance, phosphorylation, and ubiquitylation, during the first 24-hours of *Mycobacterium tuberculosis (Mtb)* infection of primary macrophages. We discovered 1379 proteins with changes in their phosphorylation state and 591 proteins with changes in their ubiquitylation in response to *Mtb* infection. We identified pathways regulated by phosphorylation and ubiquitylation that weren’t reflected by changes in protein abundance, indicating that the activity of these pathways was regulated. These include pathways known to be regulated by ubiquitylation and phosphorylation (*e.g.* autophagy) as well as pathways that were not known to be regulated during *Mtb* infection (e.g. nucleocytoplasmic transport and mRNA metabolism). We identified an enrichment in phosphorylation of autophagy receptors (TAX1BP1, p62, optineurin, BNIP3L), several of which were not previously implicated in the host response to *Mtb* infection. We found that p62 deficiency blocks ubiquitylation and TAX1BP1 deficiency enhances ubiquitylation, suggesting p62 ubiquitylation acts as an amplification loop by promoting downstream adaptor recruitment and serves as a platform for recruitment of ubiquitin. Our results show that TAX1BP1 mediates clearance of ubiquitylated *Mtb* and targets the bacteria to LC3-positive phagophores. Taken together, our proteomic profiling is likely a valuable resource for initiating mechanistic studies of macrophage biology.

## Introduction

*M. tuberculosis* infection remains a major cause of mortality from infectious disease and drug resistant isolates pose a public health threat^1–3^. Our current understanding of the mechanisms by which *Mtb* is controlled by the immune system is only partially understood. While some pathways have been identified for *Mtb* control, there are likely many other pathways that await discovery. A better understanding of the immune mechanisms that underlie resistance to *Mtb* may shed light on new therapeutic and diagnostic modalities to combat this global health threat.

One of the modalities by which macrophages sense and respond to *Mtb* is by changing transcription^4, 5^, which leads to an increased pro-inflammatory state^4, 6^. However, many other cellular processes (*e.g.* cell death pathways, autophagy, and trafficking) are not controlled by transcription^7,8^. Understanding these non-transcriptional pathways will give us a more complete picture of the ways by which macrophages attempt to control *Mtb* infection and has revealed mechanisms by which intracellular pathogens persist in host cells^9^.

Autophagy is an example of a non-transcriptional macrophage response that is controlled both by phosphorylation and ubiquitin and is thought to lead to degradation of microbes in the lysosome^10, 11^. *Mtb* perforation of the phagosomal membrane leads to activation of TANK-binding kinase 1 (TBK1), a kinase important for ubiquitin-mediated autophagy targeting as well as the induction of type I IFN^12, 13^. Some ubiquitin ligases have been identified that are required for Mtb-mediated autophagy^14, 15^, but their substrates and role is still mysterious and there are likely other ones required. Deposition of ubiquitin subsequently recruit adaptors (p62, NDP52, and Nbr1) that link the ubiquitylated cargo to autophagy by the LC3-interacting region (LIR)^16–18^. p62 has been shown to be phosphorylated and ubiquitylated in its ubiquitin binding domain, and both modifications have been shown to change the binding characteristics of the adaptor enabling recognition of polyubiquitylated cargo for selective autophagy^19, 20^. In the case of *Mtb,* some adaptors have been implicated in autophagy of the *Mtb*-containing vacuole, but not all of them have been tested.

To improve our understanding of cellular responses to infection and especially autophagy, we profiled the changes in host protein abundance, phosphorylation, and ubiquitylation during the first 24-hours of *Mtb* infection in primary macrophages. By measuring dynamic changes in phospho-serine, -threonine, and -tyrosine with a label-free approach in primary cells during a dense time course of *Mtb* infection, we expand on our previously published work profiling changes in levels of post-translational modifications at a single time point in the RAW macrophage cell line^21^, and of tyrosine phosphorylation in primary macrophages^22^. Among the over 1,300 phosphorylated peptides that changed during *Mtb* infection, we identified significant changes in phosphorylation of the autophagy receptors p62, Bnip3l, TAX1BP1, and Optineurin. We show for the first time that Optineurin and TAX1BP1 colocalize with *Mtb.* Whereas ubiquitylation of *Mtb* is diminished in p62 targeted knockout macrophages, deficiency of TAX1BP1 results in accumulation of ubiquitylated *Mtb,* and augments activation of TBK1 and recruitment of p62 to *Mtb.* In contrast, *Mtb* colocalization with LC3 is inhibited in macrophages deficient for p62 or TAX1BP1. In this study, we systematically identified thousands of post-translational modifications during macrophage infection, which enabled us to gain new insight in the distinct roles of autophagy receptors during *Mtb* infection.

## Results

### Proteome-level evaluation of primary macrophage responses to *M. tuberculosis* infection

To identify new innate immune pathways modulated during *M. tuberculosis* infection, we sought to obtain a deep data set of the changes in host protein abundance and post-translational changes during a time course of macrophage infection. To this end, we infected primary murine bone marrow-derived macrophages with *Mtb* in biological triplicate, harvested infected cells at 2-, 4-, 6-, 8-, and 24-hours post-infection, and prepared protein lysates (Fig. 1A). We also performed exact time-matched mock infections of the same macrophages and harvested at 0-, 6-, and 24-hours. Samples were digested with trypsin and a portion of the resulting peptides were set aside for abundance measurements and the remaining peptides were subjected to serial enrichment using di-gly remnant^23^ and phospho-peptide affinity technologies^24^. We subjected all peptide samples (abundance, phospho- and diGly-modified peptides) to liquid chromatography-mass spectrometry (LC-MS). Based on the measured *m/z* ratio of the parent ions and fragment ions for each peptide coupled with a database search of the mouse proteome, we determined the amino acid sequence, modification sites, and full-length protein for each peptide. As an important quality control measure, pair-wise comparison of parent ion MS intensity measurements between technical and biological replicates from the same condition were highly reproducible with robust correlation coefficients from global abundance, phosphoenriched, or diGly-enriched samples (Table S1; Fig 1B).

**Fig. 1.**
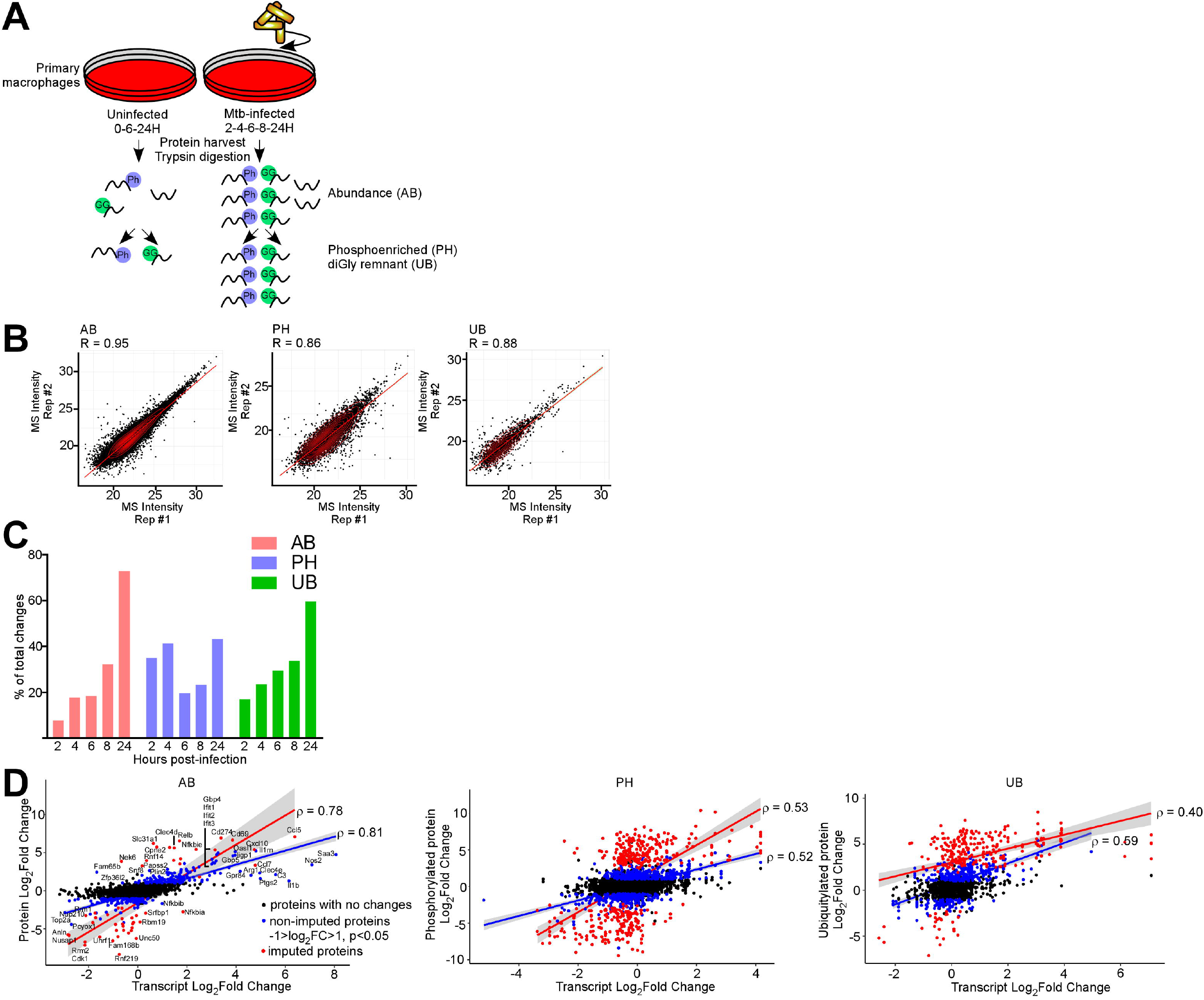
Proteomics and RNAseq profiling of *Mtb* infected bone marrow-derived macrophages. (*A*) Schematic for the experimental design indicating the hours post-infection at which point mock-infected or *Mtb* infected macrophages were harvested and peptides generated for global protein abundance measurements by LC-MS. Phosphorylated peptides (Ph) or ubiquitylated peptides containing the diGly-remnant (GG) were separately enriched. (*B*) Replica plots of MS intensity measurements for individual peptides in global protein abundance (AB), phosphopeptide (PH), or ubiquitylated peptide (UB) samples. The correlation coefficient is displayed (R). (*C*) Percent of total changes occurring at each time point 2-24-hours post-infection for global protein abundance, phosphorylation, or ubiquitylation. (*D*) Correlation plot of changes in gene transcription and protein abundance (AB), phosphorylation (PH), or ubiquitylation (UB) at 24-hours post-infection with *Mtb*. Proteins with no statistically significant changes during *Mtb* infection are colored black. Non-imputed proteins with statistically significant changes during *Mtb* infection are colored blue. Imputed proteins during *Mtb* infection are colored red. The correlation coefficient for non-imputed and imputed proteins are both shown.

To determine the statistically significant changes in protein abundance and/or PTMs in response to infection, we performed label-free quantification using MSstats to compare unfragmented (MS1) intensity peptide counts between infected and uninfected samples across the biological and technical replicates. Peptides with a log_2_-fold change in intensity of >1 or <-1 and a *p* value <0.05 were deemed statistically significant. Importantly, in many cases, we identified significant peptide values in one experimental condition (infected) but no corresponding peptides in the cognate (uninfected) control, indicating that these changes were likely biologically relevant, but which complicated the computation of a meaningful ratio. To address this issue, we used a method described by Waters *et al.* in which missing values were imputed by the limit of MS1 detection to reveal an “imputed value”^25^. An aliquot of peptide samples were analyzed for changes in global protein abundance prior to enrichment for phosphopeptides or diGly-remnant peptides (Fig. 1A). Analysis of global protein abundance, phosphorylated peptides, and di-Gly remnant peptides revealed up to 50,000 peptides matching to 4,500 unique proteins (Supplemental S1) and correlating to over 1,000 statistically significant changes in peptide levels during *Mtb* infection (Supplemental S2, S3, and S4).

Over the 24-hour time course, we observed an increase in the number of changes in abundance and ubiquitylated proteins at each subsequent time point starting at 2-hours post-infection with the maximum number detected at 24-hours post-infection (Fig. 1C). In contrast, phosphopeptide changes were bimodal with the greatest number of phosphopeptide changes occurring early (2- or 4-hours post-infection) or late (24-hours) after *Mtb* infection (Fig. 1C).

Although protein abundance levels are to a large extent determined by mRNA levels^26^, the correlation between the two is affected by the substantial delay (up to 6-hours) in protein abundance changes following changes in mRNA levels, post-translational modifications (*i.e.* ubiquitylation) affecting protein degradation, and translation rate modulation^27^. In order to determine the level of correlation between the changes in RNA and protein abundance after *Mtb* infection, we compared recently published transcriptomics dataset in murine bone marrow-derived macrophages 24-hours post-infection with *Mtb*^5^ and transcriptomics datasets from our laboratory at 2-, 6-, and 24-hours post-infection. Overall there was modest correlation between protein and RNA abundance at 24-hours post-infection for proteins using non-imputed (rho=0.81) and imputed peptide methods (rho=0.78; Fig. 1D). Weaker correlations were observed between mRNA and protein abundance changes at 2- and 6-hours post-infection (Supplemental S5), or between transcriptional changes and phosphopeptide or ubiquitylated peptide levels (Fig. 1D). These results suggest that the change in RNA abundance is a moderate predictor for change in protein abundance and that post-translational mechanisms (*i.e.* ubiquitylation targeting proteins for degradation in the proteasome) play an important role in affecting protein abundance during *Mtb* infection.

### *Mtb* infection elicits unique and overlapping changes at the translational and post-translational level

A complete list of changes in macrophage global protein abundance, phosphopeptide, and diGly-remnant peptides at 2-24-hours post-infection with *Mtb* are is found in Tables S4-S6. As an example, at 24-hours post-infection, the volcano plots highlight in green representative non-imputed peptides with statistically significant *p* values <0.05 and a log_2_-fold change >1 or <1 (Fig. 2A). Imputed peptides were assigned a *p* value just above the level of statistical significance and therefore are located immediately above the horizontal line demarcating the *p* value of 0.5 (-log_10_(p value) = 1.3; Fig 2A). Many of the proteins changing in global abundance and ubiquitylation, but to a lesser degree in their phosphorylation levels, are encoded by interferon-β and -γ regulated genes such as *Ifit1, Ifit2, Stat1,* or *Ifih1* (Fig. 2A). When comparing the individual proteins at all of the time points changing in abundance (n = 470), phosphorylation (n = 1377), or ubiquitylation (n = 591) during *Mtb* infection, we again noted that there was limited overlap between proteins that changed in global abundance and those modified at the post-translational level (Fig. 2B).

**Fig. 2.**
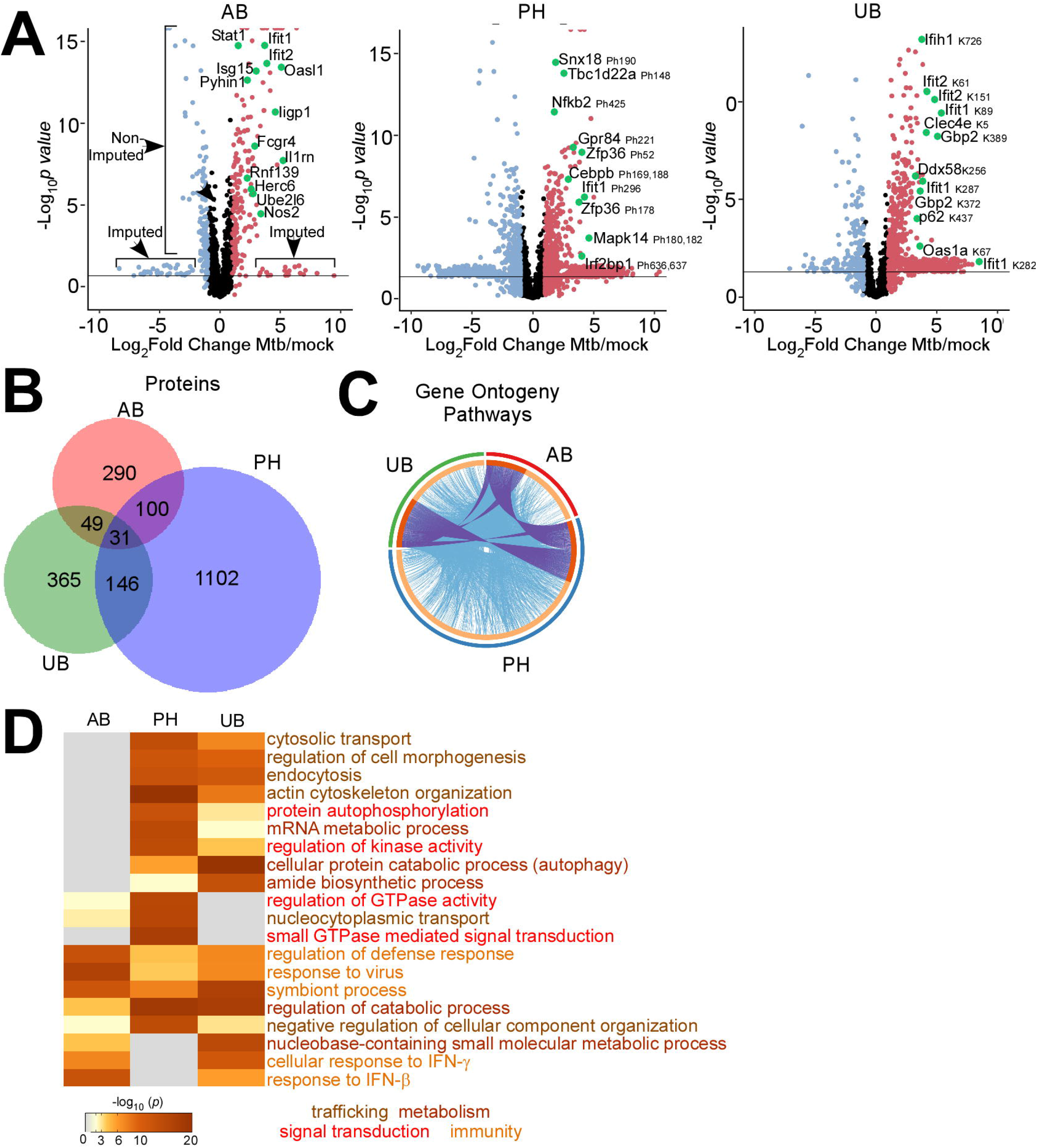
Comparison of macrophage proteins changing in abundance, phosphorylation, or ubiquitylation during *Mtb* infection. (*A*) Volcano plots displaying proteins changing in abundance, phosphorylation, or ubiquitylation at 24-hours post-infection. Proteins with a log_2_(fold change) greater than 1 are colored red. Proteins with a log_2_(fold change) less than −1 are colored blue. Proteins with a *p* value less than 0.05 (or −log_10_ (*p* value) greater than 1.3) are above the horizontal black line. Phosphorylated residues and ubiquitylated lysine residues are noted. (*B*) Venn diagram displaying the number of unique and overlapping proteins changing in abundance (AB), phosphorylation (PH), or ubiquitylation (UB) in the aggregate measurements 224-hours post-infection. (*C*) Circos visualization of protein and functional enrichment term overlap in the aggregate measurements 2-24-hours post-infection. Identical protein pairs are linked by a blue line. Protein pairs falling into the same enriched term are linked by a purple line. (*D*) Enriched ontogeny clusters highlighting commonly enriched pathways. Pathways are colored coded into common groups (trafficking, metabolism, signal transduction, immunity).

Gene ontogeny classification is one annotation method by which groups of genes or proteins are characterized by overlapping cellular function and identify pathways involved in the host responses^28^. To quantify how changes in individual protein or modification levels affect cellular pathways and gene ontogeny annotations, we performed a functional pathway meta-analysis with the web-based bioinformatics analysis pipeline, Metascape^29^. Among the 2,000 out of 9,381 gene ontogeny functional pathways enriched during infection (p<0.01), 296 were shared between global abundance, phosphorylation, and ubiquitylation enrichment groups, and each group contained hundreds of overlapping proteins or proteins with the same ontology term (Fig. 2C). The most functional pathways changing during *Mtb* infection were identified by phosphorylation (n = 1412) followed ubiquitylation (n = 1111) and global abundance (n= 719; Fig. 2C).

Enrichment meta-analysis of gene ontogeny functional pathways that changed during *Mtb* infection from peptide changes pooled at all time points revealed enrichment for immunity, cellular trafficking, metabolism, and signal transduction pathways among the top 20 most statistically significant pathways (Fig. 2D). Unique to the phosphopeptide datasets was enrichment for small GTPase mediated signal transduction, whereas cellular transport pathway enrichment was revealed by global abundance and ubiquitylation changes (Fig. 2D).

### Changes in protein abundance reveal enrichment of antiviral and inflammatory pathways during *M. tuberculosis* infection

We next sought to determine how the changes in protein abundance levels clustered together during the 24-hour time course of infection and the functional pathways represented by groups of proteins changing in the same direction following *Mtb* infection. Protein abundance and imputed protein changes each clustered into two large groups and several smaller clusters (Fig. 3A). The first group comprised of proteins increasing in abundance during *Mtb* infection and was functionally enriched for inflammatory, antiviral, and response to molecular of bacterial origin (Fig. 3B and Supplemental S6; −log_10_ *p* = 12.66, 24.60, 14.08). In contrast, the second cluster of proteins that decreased in abundance during *Mtb* and included ribosome biogenesis, DNA replication, and regulation of protein catabolism in the vacuole (Fig. 3B, Supplemental S6; −log_10_ *p* = 9.05, 10.08, 6.53). These findings are consistent with previous studies implicating a strong anti-viral response^30^ and enrichment in ribosome biogenesis pathways in transcriptional changes following *Mtb* infection^31^

**Fig. 3.**
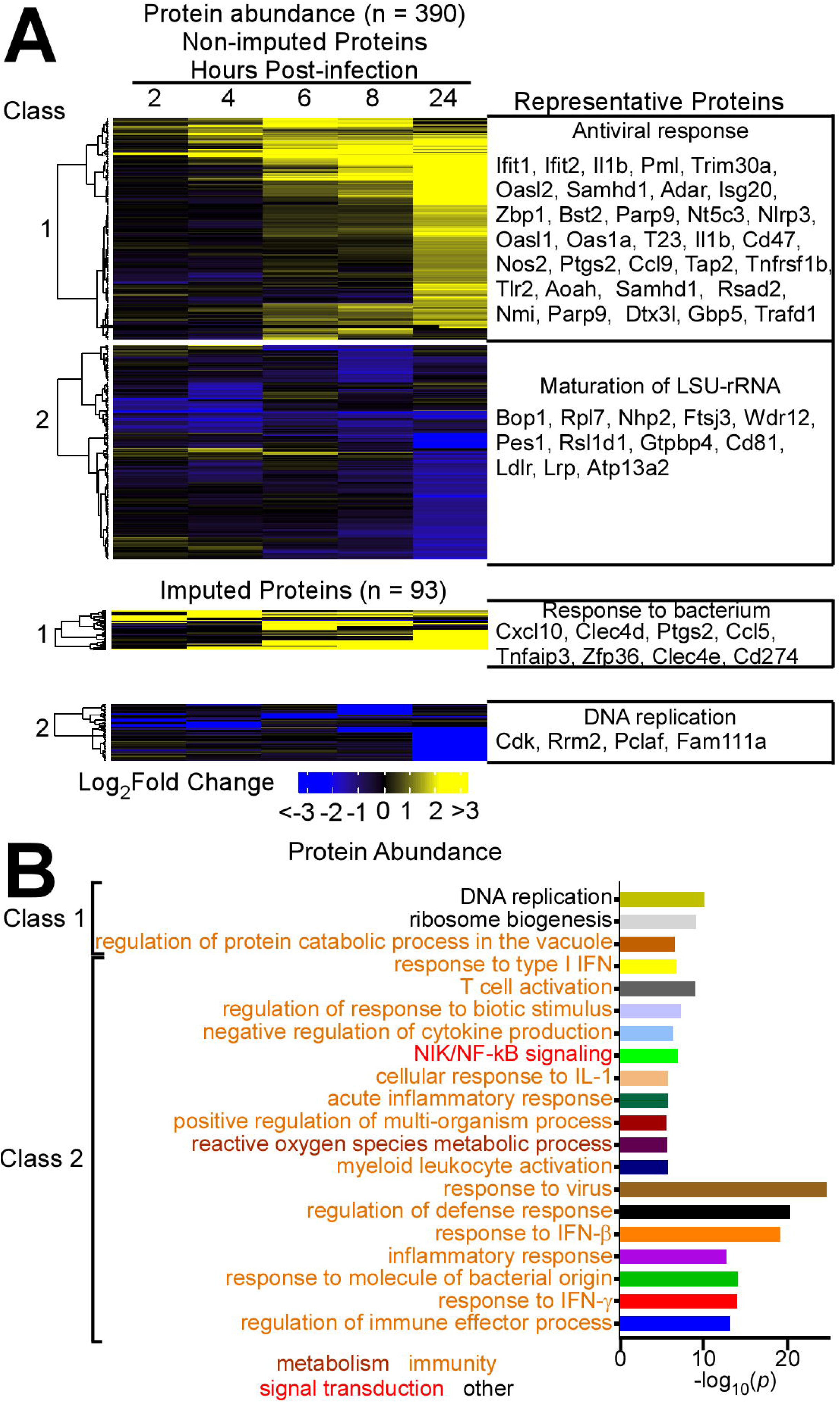
Host protein abundance changes during infection with *Mtb*. (*A*) Cluster analysis and heat map displaying the log_2_(fold change) in protein abundance for non-imputed and imputed proteins with a *p* value less than 0.05. The most statistically significant functional enrichment term and representative proteins are indicated for each class of proteins. (*B*) Graph of the functional enriched terms for protein class 1 and 2.

### Proteins ubiquitylated during *M. tuberculosis* infection enriched for immune responses and autophagy

Ubiquitylated substrates clustered together into those decreasing during infection (Cluster 1, Fig. 4A), or those increasing early (2-4-hours; Cluster 2, Fig. 4A), mid (6-8-hours; Cluster 3, Fig. 4A), or late (24-hours; Cluster 4, Fig. 4A) post-infection. While the majority of the changes in functional pathways demonstrated the most statistically significant enrichment at 24-hours such as cellular protein catabolic process, response to virus, nicotinamide nucleotide metabolism, and cellular response to interferon gamma (Fig. 4A, cluster 4; −log_10_ *p* = 13.21, 10.01, 8.30), peptide metabolic process was a notable exception, which was most enriched at 2- and 4-hours post-infection (Cluster 2, −log_10_ *p* = 11.31). Nucleosome assembly and actin cytoskeleton organization were the top two functional pathways that decreased in functional enrichment (Cluster 1, −log_10_ *p* = 6.74, 6.23; Supplemental S6). Autophagy enrichment was most statistically significant at 6- and 8-hours post-infection (Fig. 4A, cluster 3, −log_10_ *p* = 8.09). These findings are consistent with the notion that *Mtb,* in addition to eliciting an antiviral response, affects nucleosome assembly, which has been directly implicated in viral pathogenesis and HIV gene transcription during *Mtb* co-infection^32^.

**Fig. 4.**
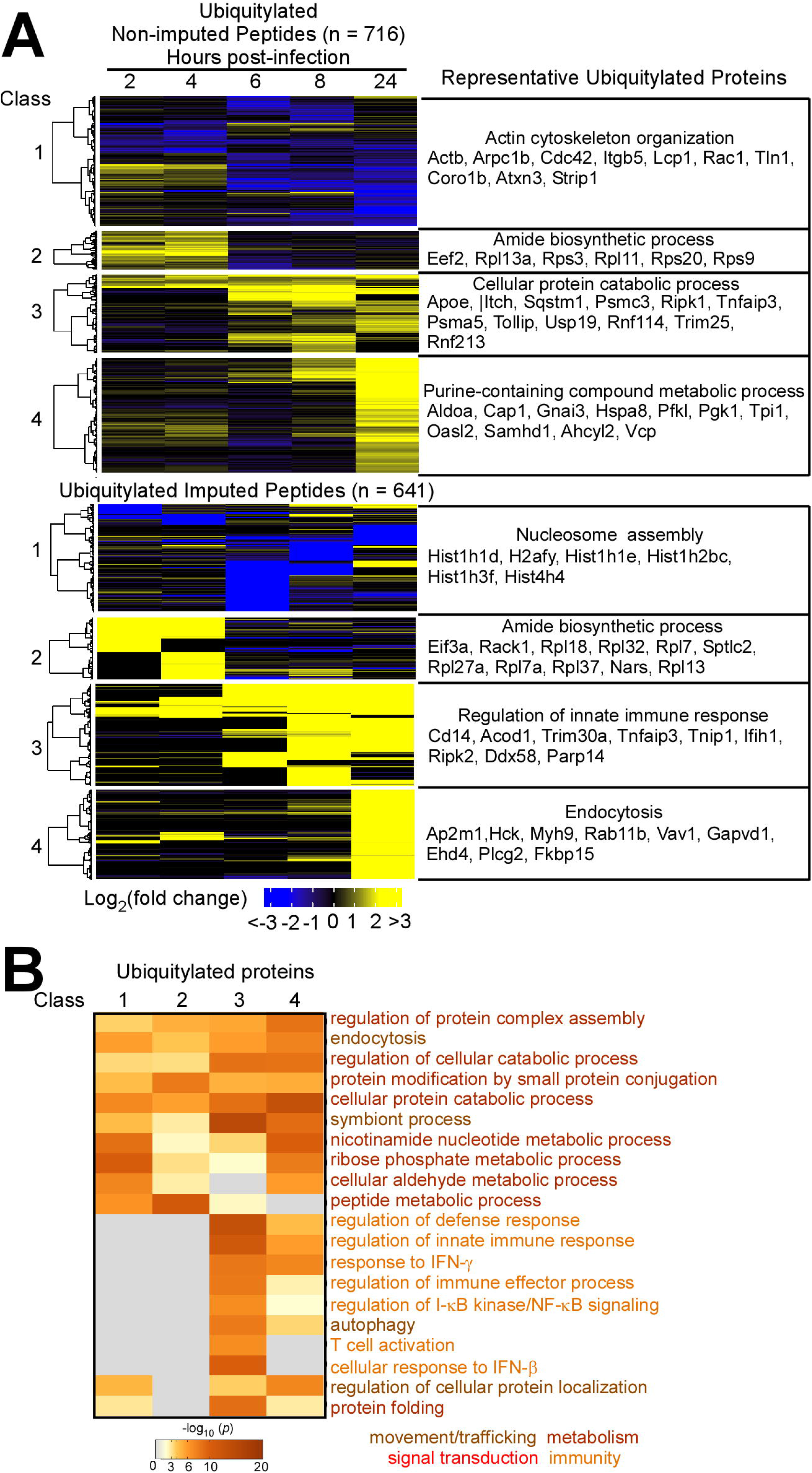
Changes in host protein ubiquitylation during *Mtb* infection. (*A*) Cluster analysis and heat map displaying the log2(fold change) in ubiquitylated peptide levels for non-imputed and imputed proteins with a *p* value less than 0.05. The most statistically significant functional enrichment term and representative proteins are indicated for each class of proteins. (*B*) Heat map visualization of functional enriched terms for each protein class.

### Changes in host protein phosphorylation during *M. tuberculosis* infection impact cellular trafficking pathways

Phosphopeptide changes during *Mtb* infection clustered into four groups: two groups increasing during infection at 2-4-hours (Fig. 5A, Class 1) or 6-24-hours (Class 2), and two groups decreasing during infection at 2-4-hours (Class 3) or 24-hours (Class 4). Functional enrichment for actin cytoskeleton organization was significant in all four clusters and especially in early in infection (Fig. 5A and B, Class 1, −log_10_ *p* = 15.29). Additional cellular trafficking pathways changing during *Mtb* infection included endosomal, cytosolic, and nuclear transport (Fig. 5B). *Mtb* engaged signal transduction components including small GTPase and I-κB kinase/NF-κB (Cluster 3, −log_10_ *p* = 11.95 and 9.29). This data suggests a prominent role for cellular trafficking pathways in the host response to *Mtb* infection at all of the measured time points and highlights the inflammatory signaling molecules activated during *Mtb* infection.

**Fig. 5.**
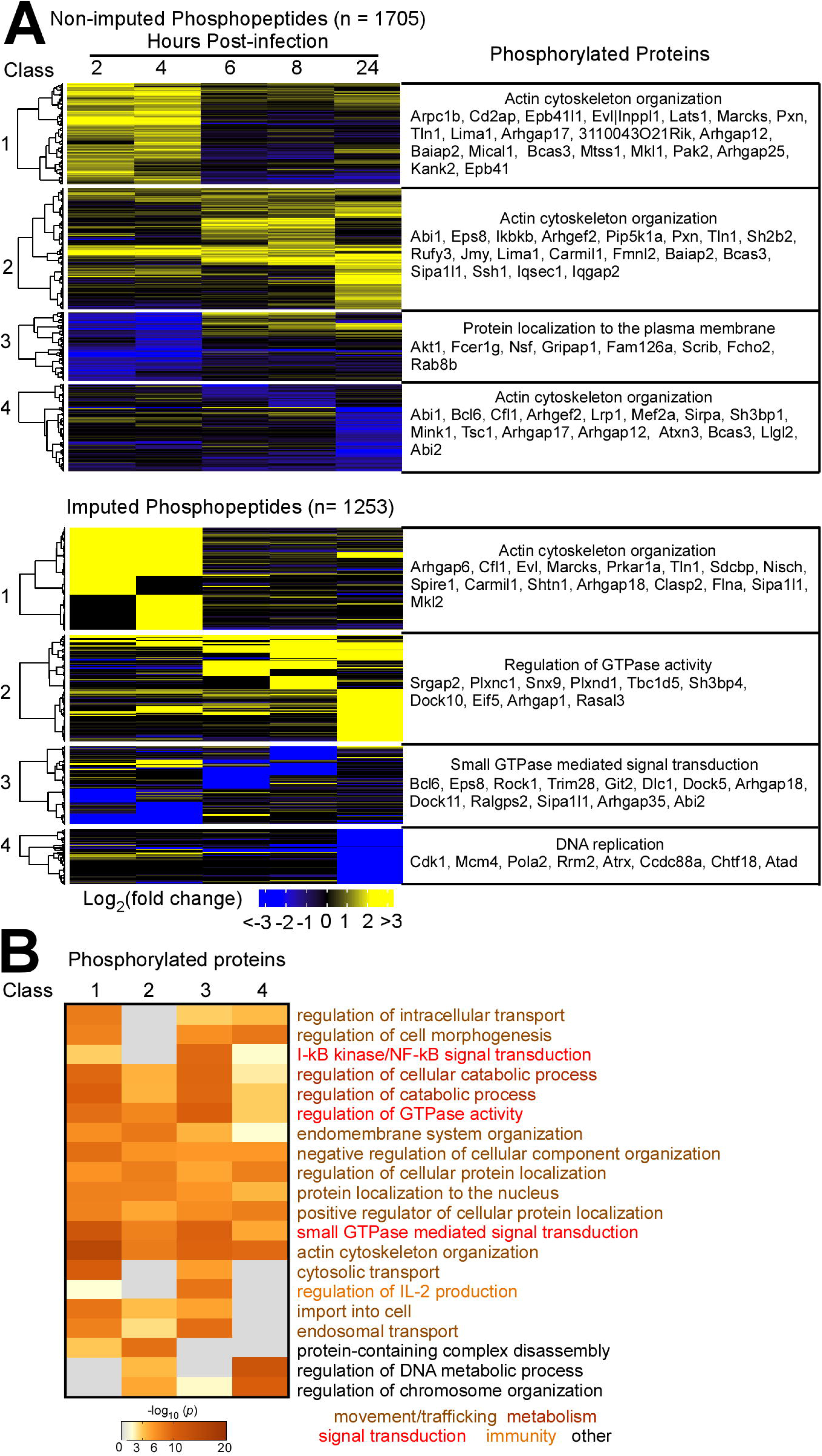
Changes in host protein phosphorylation during *Mtb* infection. (*A*) Cluster analysis and heat map displaying the log2(fold change) in phosphorylated peptide levels for non-imputed and imputed proteins with a *p* value less than 0.05. The most statistically significant functional enrichment term and representative proteins are indicated for each class of proteins. (*B*) Heat map visualization of functional enriched terms for each protein class.

### Kinase prediction tools identified kinase families and ubiquitin ligase complexes changing during *M. tuberculosis* infection

Although the functional significance of these phosphorylation changes await determination, bioinformatic tools that rely on the extremely large number of phosphoproteomic studies performed to date, allow for prioritization of those modification that likely regulate protein function and can infer when a specific kinase is regulated^33^. The large number of phosphorylation changes, and the temporal specificity, indicates that many kinases are activated during the 24-hour interaction of macrophages with *Mtb.* To begin to identify these kinases, we used a bioinformatics approach to compare the substrates phosphorylated during *Mtb* infection with a database of known kinase substrate pairs. Interestingly, several potential kinases were identified from the earliest phosphorylation data, including GTF2F1, Aurora, and ITR, suggesting that these may be good targets for prioritization of signaling cascades activated early upon interaction. Indeed, Aurora kinase is interesting given its recently discovered relationship with HIV infection, and it has not been previous implicated in bacterial infection^34^. We also sought to predict the kinases involved in trafficking and motility (Fig. 6). Indeed, we predicted the activity of important kinases that regulate actin filament dynamics, rho-associated protein kinase 1 and its kinase substrates including LIMK1 and LIMK2, changed during the time course of *Mtb* infection (Fig. 6A). Consistent with the canonical role in JAK-STAT mediated activation of immune cell survival, activation, and recruitment and MAP2Ks in the stress-response, the kinase prediction tool revealed increased kinase activity of JAK as early as 2-hours post-infection and maximal levels of MAP2K activity at 24-hours post-infection (Fig. 6A). IKB kinase, another key inflammatory kinase involved in NF-κB signaling^35^, was also predicted to undergo change in its kinase activity during *Mtb* infection.

**Fig. 6.**
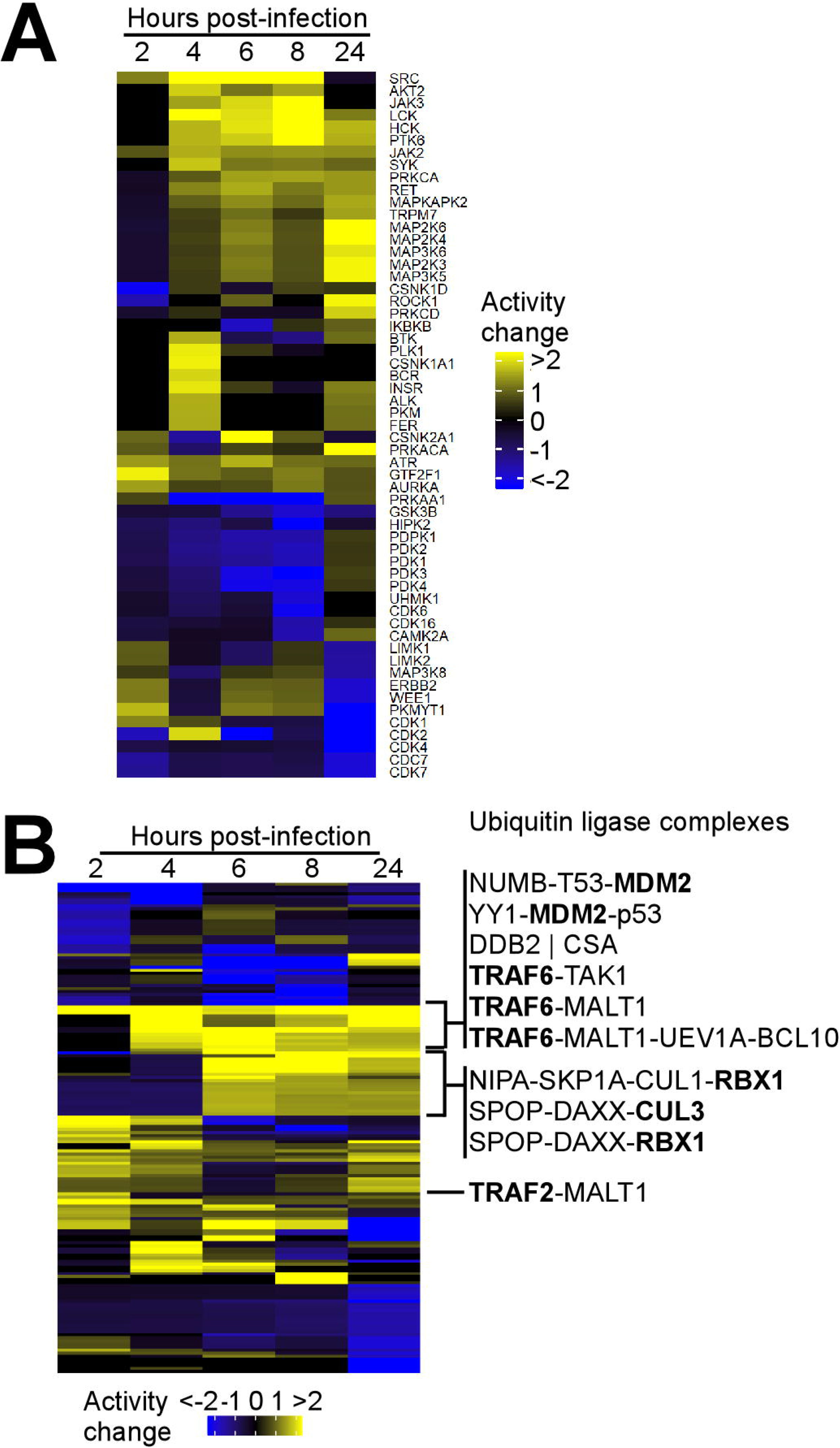
Protein kinase and effector complex activity predictions. (*A*) Cluster analysis and heat map displaying the predicted host protein kinase activity. Mouse phosphopeptide sites were mapped to the human proteome and kinase activity predictions were performed with PhosFate^33^. (*B*). Effector complex activity predictions. Effector complexes containing ubiquitin ligases (bold) are listed.

Similarly, potential ubiquitin ligases can be prioritized by their presence in kinase complexes based on a bioinformatic search of the predicted kinase complexes from our phosphopeptide datasets. In addition to revealing the TRAF-2 and −6 predicted ubiquitin ligase complexes, which have been previously implicated in immune signaling during infection^36^, the prediction tool identified previously unrecognized predicted ubiquitin ligase complexes including the ligases CUL3, RBX1, and MDM2 (Fig. 6B).

### p62, NBR1, optineurin, and TAX1BP1 colocalize with *M. tuberculosis* during macrophage infection

The autophagy pathway was prominently enriched in both PTM datasets, and we noted a significant number of phosphorylation events that occurred on autophagy adaptors. Indeed, our results identified many of the previously recognized phosphosites, including Ser-152 in the RIPK1 domain and Ser-226 in the TRAF6 binding domain of p62^19^, but we have identified novel residues as well. Importantly, as our understanding of all the adaptors that participate in targeting *Mtb* is incomplete, we noted several autophagy adaptors that were not previously implicated in the response were phosphorylated during infection. Among the cellular networks formed by the phosphorylated proteins during *Mtb* infection, we noted a cluster containing the autophagy receptors p62, optineurin, and TAX1BP1 (Fig. 7A). While p62 was previously shown to target *Mtb* to the autophagosome^16^, the other adaptors with changes in their phosphorylation levels had hitherto not been implicated in autophagy targeting of *Mtb.* Autophagy receptor phosphorylation occurred in functional domains of the receptors (Fig. 7B) and changed throughout the time course of *Mtb* infection (Fig. 7C). Of note, the phosphosite we detected in TAX1BP1 is one of the two previously reported serine residues phosphorylated by IKKα in response to TNFα and IL-1, which results in downregulation of pro-inflammatory NF-κB through assembly of the A20 complex^37^. To test the functional consequences of the autophagy receptors phosphorylated during *Mtb* infection, we engineered FLAG-tagged constructs of seven autophagy receptors and transduced RAW macrophages with lentiviral expression vectors for each plasmid. Immunoblotting of transduced macrophages with a FLAG antibody revealed the presence of epitope tagged protein correlating to the predicted protein mobility on SDS-PAGE (Fig. 8A)^38^. We detected unique bands at molecular weight of 92 kDa and 120 kDa consistent with previously published mobilities for immunoblots of TAX1BP1 and NBR1 (Fig. 8B)^39, 40^.

**Fig. 7.**
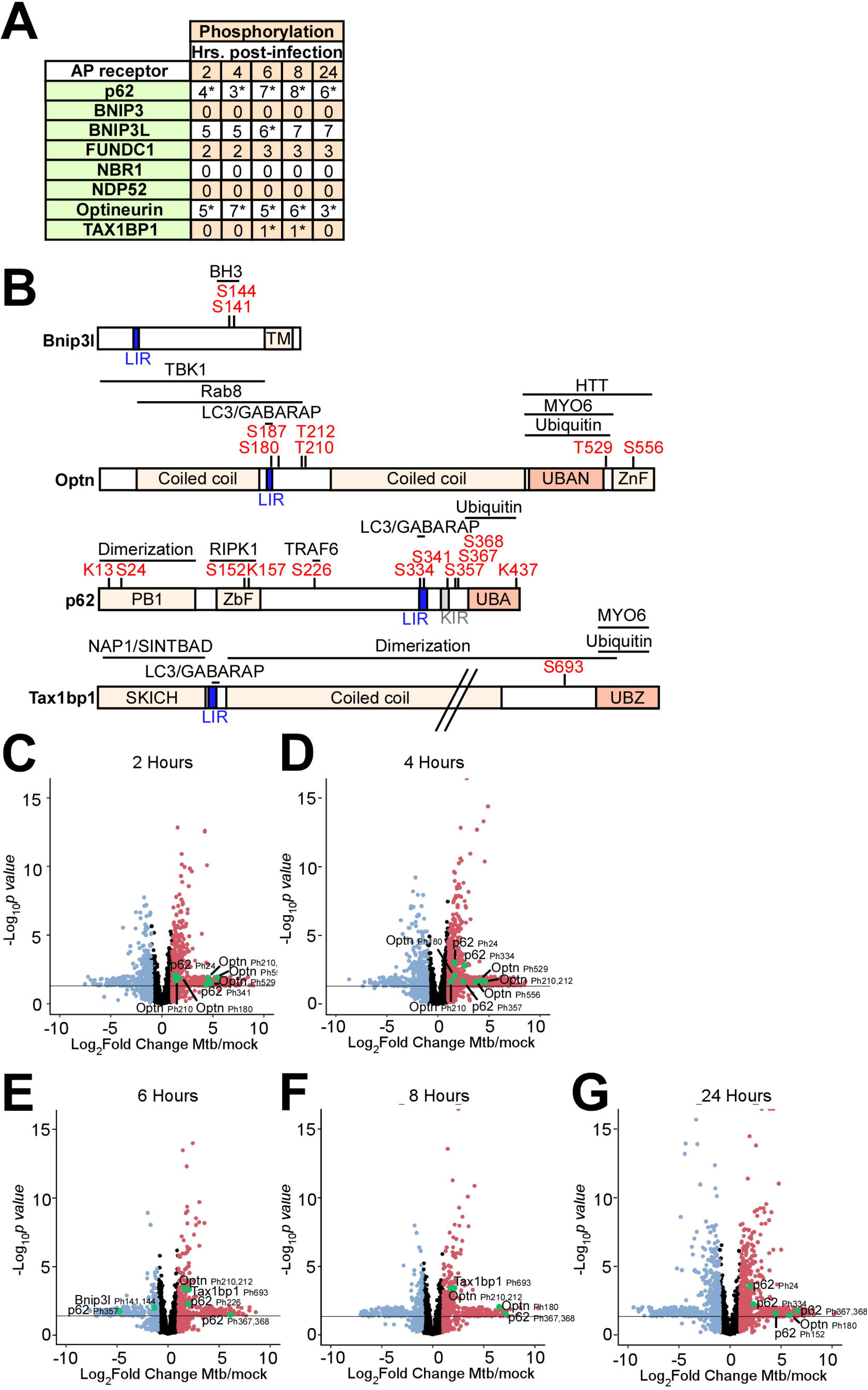
Autophagy receptor phosphorylation during *Mtb* infection. (*A*) Table showing the number of unique phosphosites in each autophagy receptor at 2-24-hours post-infection. * *p* value less than 0.05 and log_2_(fold change) greater than 1 or less than −1 for at least one of the phosphosites. (*B*) Domain organization displaying post-translational modifications in autophagy receptors. The LC3-interacting region (LIR), ubiquitin binding domain (UBAN, UBZ), myosin-6 binding domain (MYO6), SKIP carboxyl homology domain (SKICH), and TANK-binding kinase-1 (TBK1) binding domains are labeled. (*C-G*) Volcano plots highlighting changes in autophagy receptor phosphorylation at each time point 2-24-hours post*-Mtb* infection. Proteins with a log_2_(fold change) greater than 1 are colored red. Proteins with a log_2_(fold change) less than −1 are colored blue. Proteins with a *p* value less than 0.05 (or −log_10_ (*p* value) greater than 1.3) are above the horizontal black line. Phosphorylated residues and ubiquitylated lysine residues are noted.

**Fig. 8.**
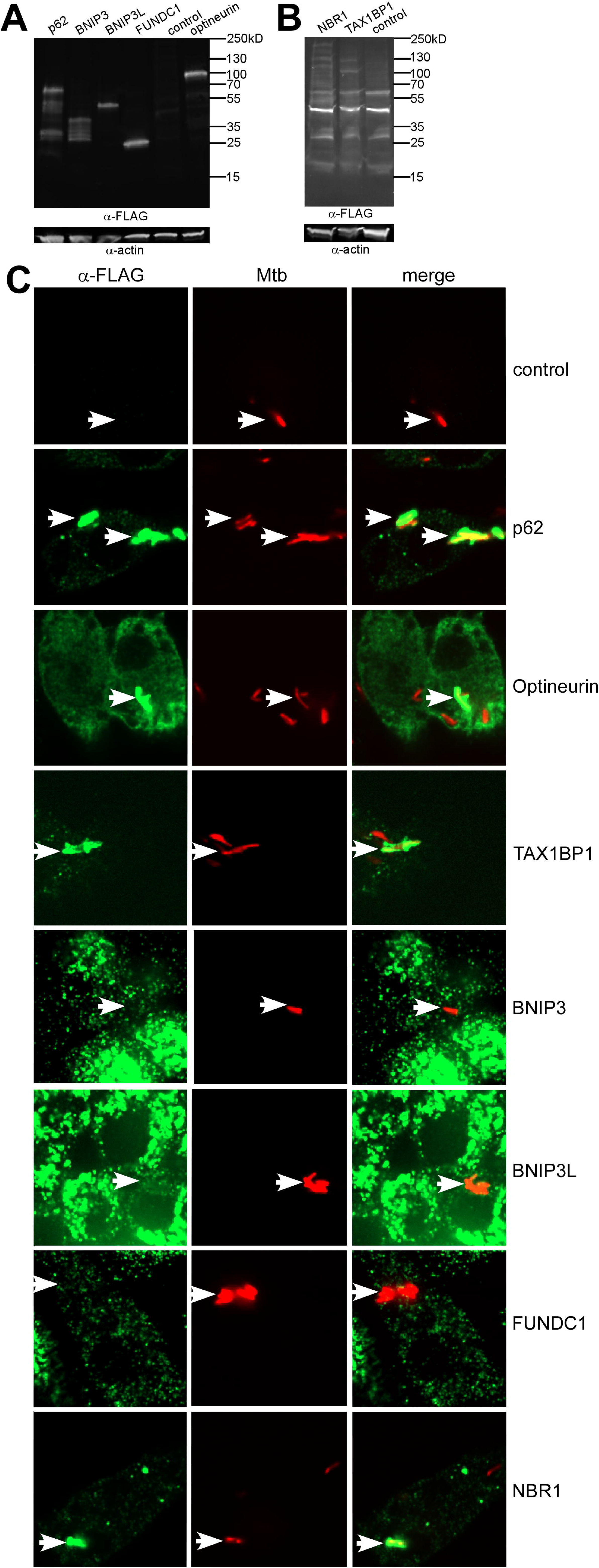
TAX1BP1, p62, optineurin, and NBR1 colocalize with *Mtb. (A* and *B*) Immunoblot showing expression of FLAG-tagged autophagy receptors in RAW macrophages. RAW macrophages were transduced with FLAG-tagged autophagy receptor constructs and cell lysates probed with α-FLAG and α-actin antisera. Immunoblots were imaged with a short exposure time (*A*) or a long exposure time (*B). (C*) Fluorescence images of RAW macrophages expressing FLAG-tagged autophagy receptors (green) with mCherry-expressing *Mtb* (red).

To directly assess for recruitment of autophagy receptors to *Mtb* during macrophage infection, we performed immunofluorescence microscopy using antibodies that recognize the FLAG epitope at 6-hours post-infection (Fig. 8C). Consistent with previous results^16^, 33%±1.7% (SEM) of mCherry-expressing *Mtb* colocalized with p62. We also detected co-localization of *Mtb* with NBR1 (29%±3.3% co-localization), Optineurin (7%±1%), and TAXBP1 (26%±3.2%). In contrast, a diffuse staining pattern without *Mtb* colocalization was observed in macrophages expressing BNIP3, BNIP3L, or FUNDC1. Together, these results revealed that p62, Optineurin, and TAX1BP1 are phosphorylated during *Mtb* infection and also colocalize with *Mtb.*

### TAX1BP1 deficiency results in accumulation of ubiquitylated *M. tuberculosis*

We used a genetic approach to evaluate the contribution of individual autophagy receptors in the selective autophagy of *Mtb.* Previous data showed deficiency of the autophagy receptors p62, NBR1, and NDP52 abrogated targeting of *Mtb* to autophagosomes^15, 41^. In contrast, loss of TAX1BP1 caused an accumulation of ubiquitin-positive *Salmonella* in a manner requiring its myosin-VI binding domain, implicating a critical role for TAX1BP1 in mediating downstream steps of autophagy targeting. To test the hypothesis that deficiency of autophagy receptors impacts the feedback loop by which autophagy kinases (*i.e.* phosphorylated TBK1), autophagy adaptors, and ubiquitin assemble around phagocytosed microbes^42^, we performed high throughput immunofluorescence microscopy and automated colocalization analysis to quantify colocalization of these autophagy markers with mCherry-expressing *Mtb* in infected *p62^−/−^* or *Taxlbp1^−/−^* targeted knockout bone marrow-derived macrophages (Fig. 9A). Consistent with prior results showing that p62 deficient macrophages are impaired in the targeting of *Salmonella* to ubiquitin^43^, we found decreased targeting of *Mtb* to ubiquitin (FK2) in *p62^−/−^* macrophages. In contrast, infected *Tax1bp1^−/−^* cells displayed increased *Mtb* colocalization with ubiquitin, phospho-TBK1, and p62 (Fig. 9B). Together these findings are consistent with a role for p62 in signal amplification of cargo ubiquitylation and TAX1BP1 in targeting intracellular pathogens including *Salmonella^40^* and *Mtb* to fully mature autophagosomes.

**Fig. 9.**
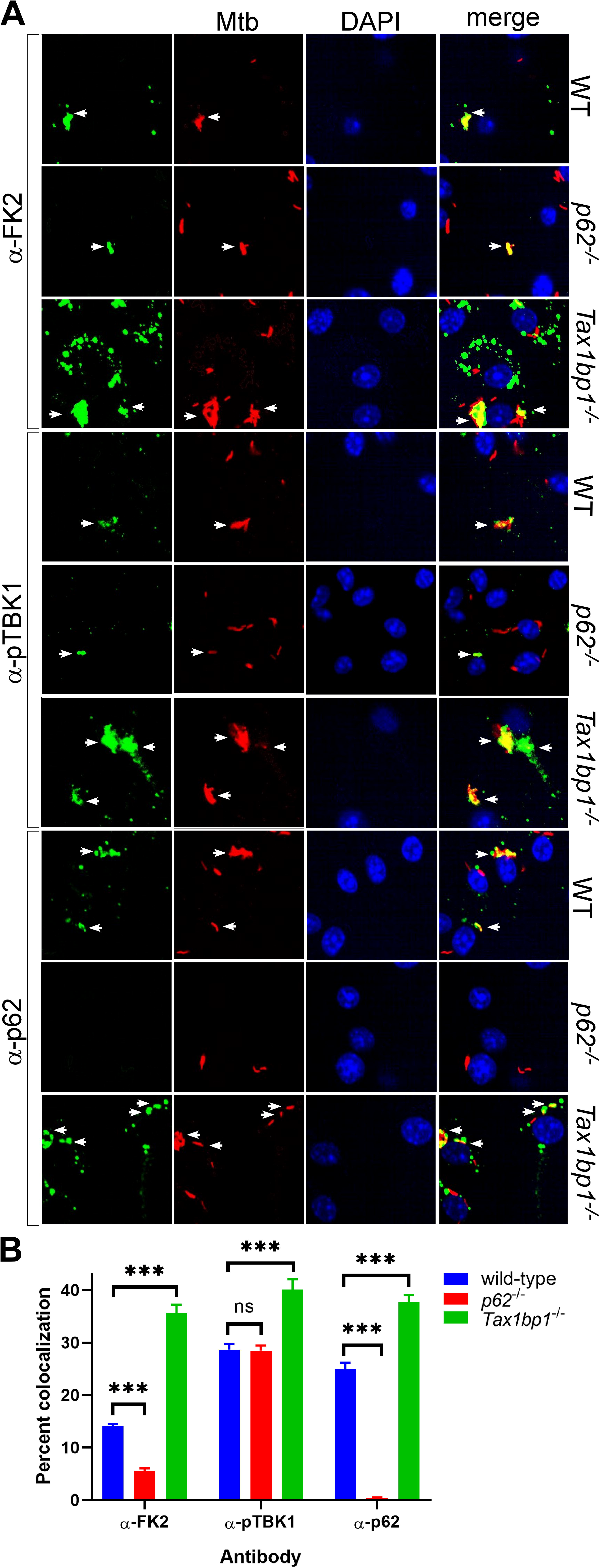
Accumulation of *Mtb* targeted to ubiquitin (FK2), p62, and phospho-TBK1 in TAX1BP1 targeted knockout macrophages. (*A*) Fluorescence images of wild-type, *p62^−/−^*, or *Tax1bp1^−/−^* bone marrow-derived macrophages infected with mCherry-expressing *Mtb* (red) and immunostained with antibodies to ubiquitin (FK2), p62, or phospho-TBK1 (green). (*B*) Quantitative analysis of *Mtb* colocalization with ubiquitin, p62, and phospho-TBK1 and 8-hours post-infection. Results are the means ±SEM for five technical replicates and representative of three independent experiments. *** *p* value less than 0.001 by t-test. ns (non-significant).

We next sought to evaluate by an alternative approach whether autophagy components accumulate around *Mtb* in TAX1BP1-deficient macrophages. We generated Cas9-expressing bone-marrow derived macrophages transduced with guide RNAs targeting *Tax1bp1* 9 or 10. In pooled populations of edited macrophages, editing efficiency at exon 9 or 10 calculated by TIDE analysis^44^ was 90.0% or 84.8%, respectively. Compared to macrophages transduced with scramble control guides, infected *Tax1bp1* edited macrophages displayed increased *Mtb* colocalization with ubiquitin, phospho-TBK1, and p62 (Fig. 10). However, infected *Tax1bp1* edited or deficient macrophages displayed decreased *Mtb* colocalization with LC3 (Fig. 11). Thus, TAX1BP1 plays an important role in the clearance of ubiquitin-positive Mtb, recruitment of p62 and phospho-TBK1, and delivery of *Mtb* to LC3. In summary, our model is that TAX1BP1 is necessary for complete targeting of *Mtb* to the autophagosome in a step downstream from recruitment of p62 and ubiquitin (Fig. 12).

**Fig. 10.**
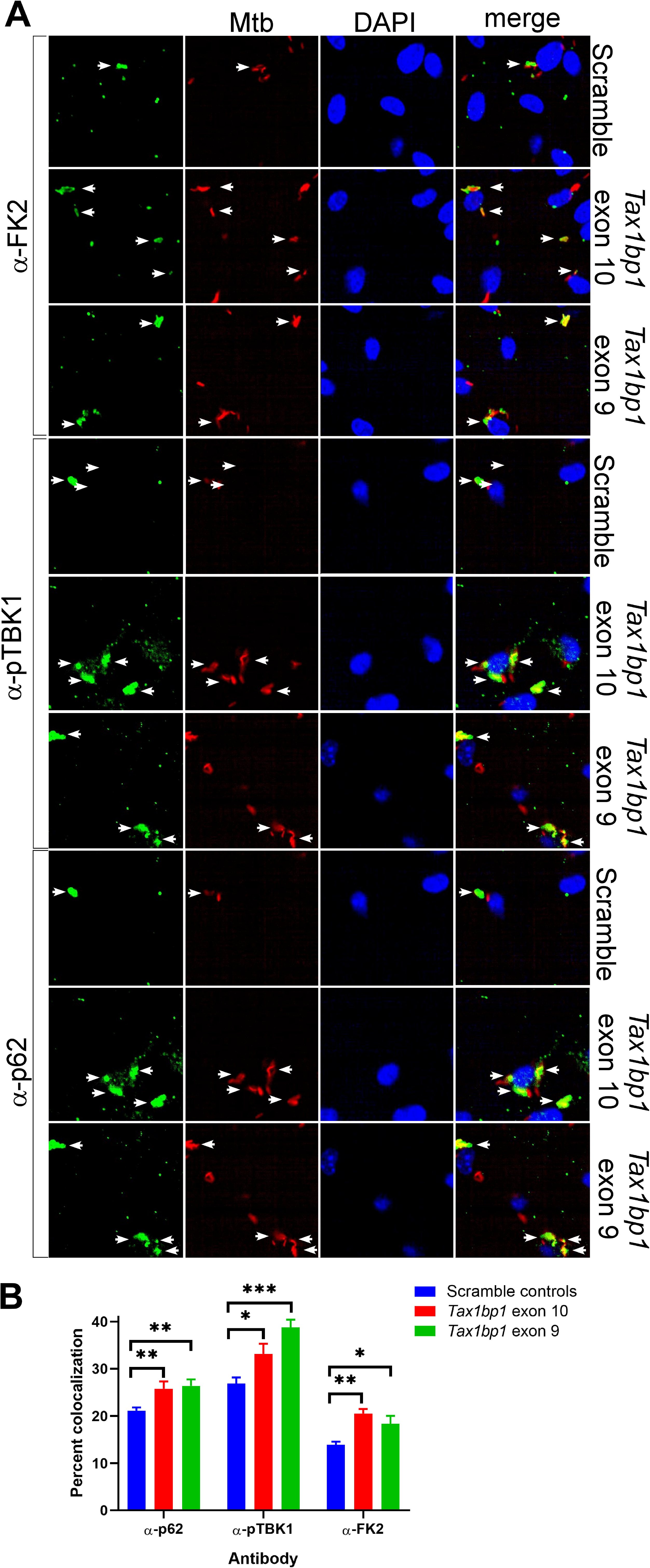
Accumulation of *Mtb* targeted to ubiquitin (FK2), p62, and phospho-TBK1 in TAX1BP1 edited macrophages. (*A*) Fluorescence images of bone marrow-derived macrophages transduced with scramble control guide RNAs or *Tax1bp1* exon 10 or 9 guide RNAs infected with mCherry-expressing *Mtb* (red) and immunostained with antibodies to ubiquitin (FK2), p62, or phospho-TBK1 (green). Nuclei were stained with DAPI (blue). (*B*) Quantitative analysis of *Mtb* colocalization with ubiquitin, p62, and phospho-TBK1 at 8-hours post-infection. Results are the means ±SEM for four technical replicates. * *p* value less than 0.05 by t-test. ** *p* value less than 0.01. *** *p* value less than 0.001.

**Fig. 11.**
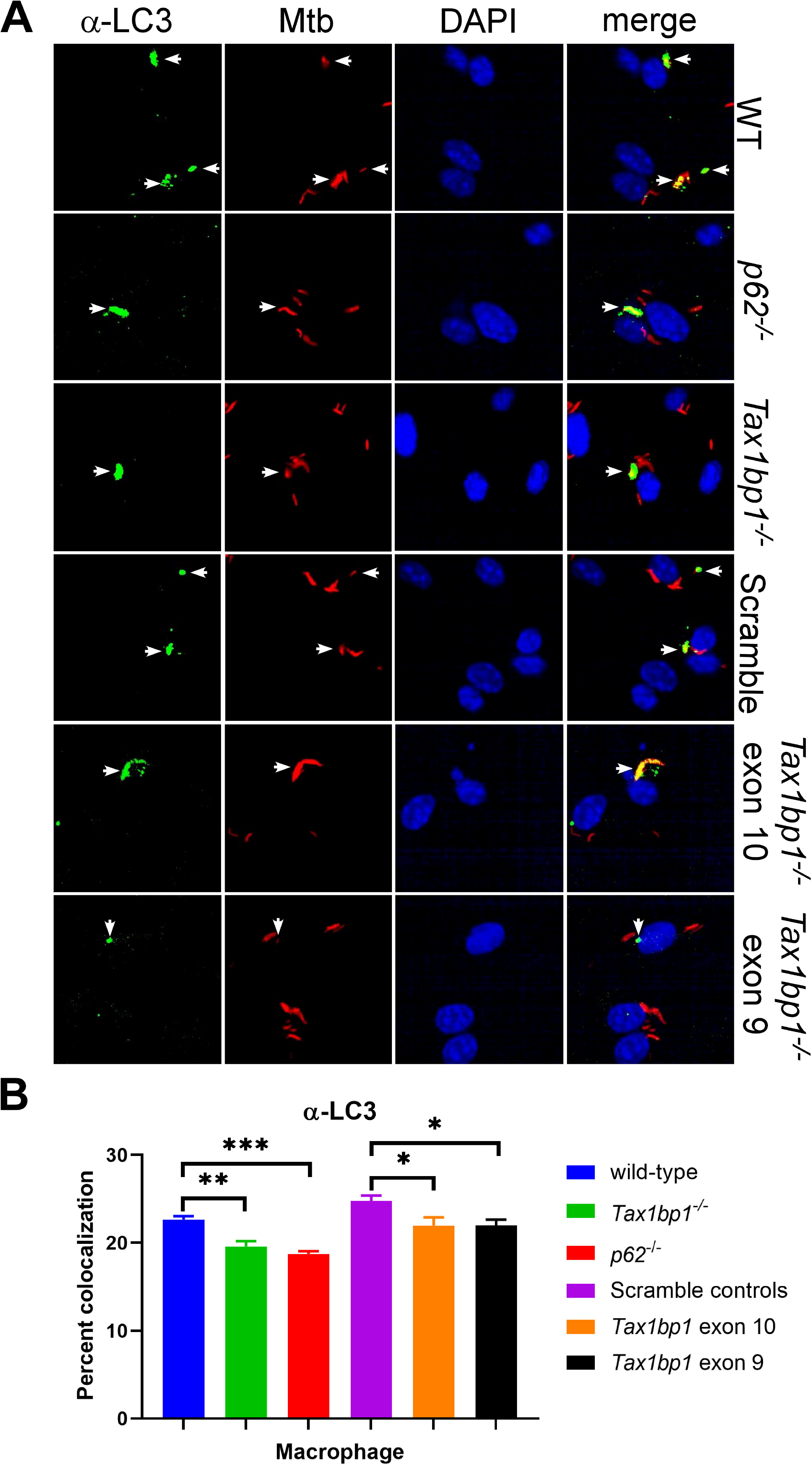
TAX1BP1 deficiency abrogates targeting of *Mtb* to LC3. (*A*) Fluorescence images of bone-marrow derived macrophages infected with mCherry-expressing *Mtb* (red) and immunostained with α-LC3 antisera (green). Nuclei were stained with DAPI (blue). (*B*) Quantitative analysis of *Mtb* colocalization with LC3 at 8-hours post-infection. Results are the means ±SEM for four technical replicates and representative of two independent experiments. * *p* value less than 0.05 by t-test. ** *p* value less than 0.01. *** *p* value less than 0.001.

**Fig. 12.**
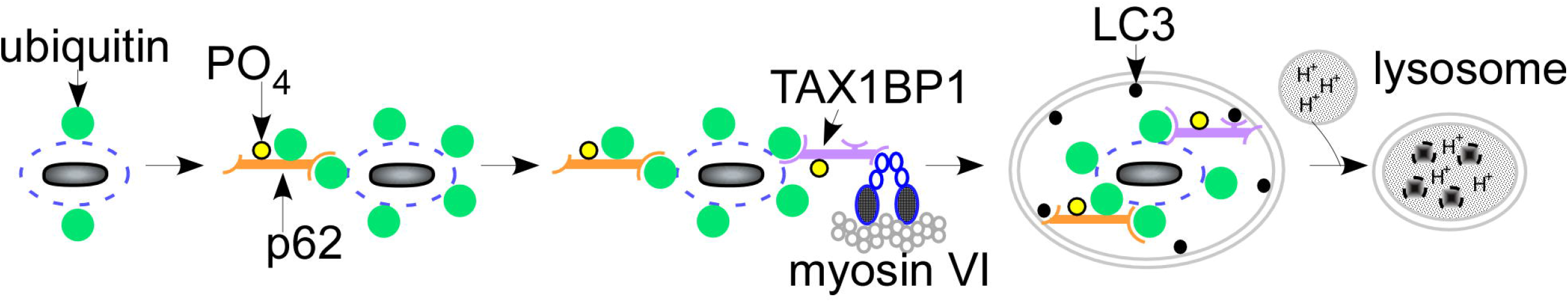
A model showing the role of p62 in amplification of ubiquitin recruitment to *Mtb* and TAX1BP1 in completing the targeting of *Mtb* to selective autophagy in a step downstream of p62 recruitment.

## Discussion

Despite the global impact of *Mtb* infection on humanity, relatively little is known about the signaling events that occur in the innate immune cells to first encounter this pathogen during infection. Phosphorylation mediated by the serine/threonine kinase, TBK1^13^, and ubiquitylation^14,15^ are two important signaling mechanisms that affect *Mtb* survival, autophagy targeting, and inflammatory responses. However, the substrates phosphorylated or ubiquitinylated during *Mtb* infection were not well described. We therefore set out to comprehensively measure the changes in host protein phosphorylation, ubiquitylation, and abundance, in primary macrophages at five time points following *Mtb* infection. This led to the identification of TAX1BP1, an autophagy receptor not previously known to be involved in *Mtb* infection. With a genetic approach, we found that TAX1BP1 is important for clearance of ubiquitylated *Mtb* and autophagosome maturation. Further work will be necessary to evaluate the impact of TAX1BP1 on *Mtb* growth and inflammatory responses, as well as the functional consequences of the thousands of other substrates that changed in their global abundance or PTM levels during *Mtb* infection.

TAX1BP1 appears to play a distinct role in antibacterial autophagy. Intriguingly, TAX1BP1 is not required for global autophagy in T cells or MEFs^45^, indicating that TAX1BP1 may play a role in targeting select cargos. However, TAX1BP1 has been implicated as an autophagy adaptor important for clearance of *Salmonella* in a manner that requires myosin VI, one of TAX1BP1’s binding partners^40^. There is also evidence that TAX1BP1 plays a role in the regulation of inflammatory responses as it is critical for the termination of inflammatory responses triggered by NF-κB signaling^46^, and binds to the pro-inflammatory tumor necrosis factor receptor associated factor-6 (TRAF6)^47^ and the anti-inflammatory A20 ubiquitin editing enzyme^48^. Likewise, the fact that it binds to myosin VI indicates that it may play a role in physical movement of large cargo within the cell perhaps to facilitate fusion with the lysosome. Given that TAX1BP1 regulates several different innate immune responses, we aim to determine how TAX1BP1 deficiency ultimately affects *in vivo* control of *Mtb* infection.

## Methods

### Ethics statement

An animal use protocol (AUP-2015-11-8096) for mouse use was approved by the Office of Laboratory and Animal Care at the University of California, Berkeley, in adherence with guidelines from the Guide for the Care and Use of Laboratory Animals of the National Institutes of Health.

### Mice and Macrophages

Wild-type C57BL/6J mice obtained from Jackson laboratories. *p62^−/−^* mice were provided by Herbert Virgin at Washington University, St. Louis, USA. *Tax1bp1^−/−^* mice were obtained from Hidekatsu Iha at the University of Oita, Japan. Rosa26-Cas9 knock-in mice^49^ were provided by Gregory Barton at the University of California, Berkeley, USA. Primary murine bone-marrow derived macrophages (BMDMs) were prepared by flushing femurs from 8-12-week-old male and female mice. Bone marrow extracts were differentiated for 7 days and cultured in DMEM-H21 supplemented with 20% FBS and 15% MCSF derived from 3T3-MCSF cells. Raw264.7 cells were obtained from the ATCC and cultured in DMEM-H21 supplemented with 10% FBS and 20 mM HEPES.

### Bacterial strain

*M. tuberculosis* (Erdman) was used for macrophage infections. *M. tuberculosis* was grown to log phase in 7H9 liquid media (BD) supplemented with 10% Middlebrook OADC (Sigma), 0.5% glycerol, 0.05% Tween80 in roller bottles at 37°C.

### Macrophage infection with *M. tuberculosis*

For proteomics, three independent experiments were performed. BMDMs were seeded at a density of 3 × 10^6^ cells per well in 4-well cell culture treated dishes. Six dishes were seeded for each experimental condition. The cells were cultured at 37°C with 5% CO_2_ for 72-hours prior to infection.

For microscopy to detect FLAG-tagged autophagy receptor colocalization, three coverslips for each experimental condition were seeded with RAW macrophages at a density of 1.2 × 10^5^ cells per coverslip in a 24-well dish. For microscopy to detect ubiquitin, p62, and phospho-TBK1, four wells for each experimental condition were seeded with bone-marrow derived macrophages at a density of 3 x 10^4^ cells per well in a 96-well plate. The cells were cultured for 18-hours prior to infection.

In order to synchronize the start time of the infections, macrophages were infected with *M. tuberculosis* (Erdman strain) using a ‘spinfection protocol’ at a multiplicity of infection of 10 for proteomics or 2 for microscopy experiments. Logarithmic phase *M. tuberculosis* suspensions were washed twice with PBS followed by centrifugation for 5 minutes at 183g and sonication to remove and disperse clumps, respectively. *M. tuberculosis* was resuspended in DMEM with 10% horse serum. Media was removed from the macrophage monolayers, the bacterial suspension was overlaid, and centrifugation was performed for 10 minutes at 183g. For proteomics experiments, cells were returned to macrophage media and incubated until harvesting at 2-, 4-, 6, 8-, or 24-hours post-infection. Uninfected mock controls were harvested at 0-, 6-, and 24-hours intervals. For microscopy experiments, infected monolayers were washed twice with PBS and then macrophage media was overlaid. At 8-hours post-infection, the macrophages were washed once with PBS, fixed with 4% paraformaldehyde, washed twice with PBS, and immunostained.

### Cell lysis

At each time point 0-24-hours post-infection, macrophages were washed with PBS warmed to 37°C and *M. tuberculosis* was sterilized on the cell culture plates by the addition of 100% methanol at 4°C. The cells were washed twice with PBS and lysed with 6 ml of lysis buffer (8 M urea, 150 mM NaCl, 100 mM ammonium bicarbonate, pH 8; added per 10 ml of buffer: 1 tablet of Roche mini-complete protease inhibitor EDTA free and 1 tablet of Roche PhosSTOP tablet) prepared fresh before each replicate. Lysates were stored at −80°C until further processing.

### Trypsin digest and desalting

Lysates were sonicated three times with a Sonics VibraCell probe tip sonicator at 7 watts for 10 seconds. In order to remove insoluble precipitate, lysates were centrifuged at 16,100g at 4°C for 30 minutes. A bicinchoninic acid assay (Pierce) was performed to measure protein concentration in cell lysate supernatants. 10 mg of each clarified lysate was reduced with 4 mM tris(2-carboxyethyl)phosphine for 30 minutes at room temperature and alkylated with 2 mM iodoacetamide for 30 minutes at room temperature in the dark. Remaining alkylated agent was quenched with 10 mM 1,4-dithiothreitol for 30 minutes at room temperature in the dark. The samples were diluted with three starting volumes of 100 mM ammonium bicarbonate, pH 8.0, to reduce the urea concentration. Samples were incubated with 100 μg of sequencing grade modified trypsin and incubated at room temperature with rotation for 20 hours. The sample pH was reduced to approximately 2.0 by the addition of 10% trifluoroacetic acid (TFA) to a final concentration of 0.3% trifluoroacetic acid followed by 6 M HCl at a 1:100 volume ratio. Insoluble material was removed by centrifugation at 1989g for 10 minutes.

Peptides were desalted using SepPak C18 solid-phase extraction cartridges (Waters). The columns were activated with 1 ml of 80% acetonitrile (ACN), 0.1% TFA, and equilibrated with 3 ml of 0.1% TFA. Peptide samples were applied to the columns, and the columns were washed with 3 ml of 0.1% TFA. Peptides were eluted with 1.1 ml of 40% ACN, 0.1% TFA. Peptides were divided for global protein analysis (10 μg), phosphoenrichment (1 mg), or diGly-enrichment (remaining sample).

### Global protein analysis

10 μg of peptides were dried with a centrifugal evaporator and stored at −20°C until analysis by liquid chromatograph and mass spectrometry.

### Phosphopeptide enrichment by immobilized metal affinity chromatography

Iron nitriloacetic acid (NTA) resin were prepared in-house by stripping metal ions from nickel nitroloacetic acid agarose resin with 500 mM ethylenediaminetetraacetic acid, pH 8.0 three times. Resin was washed twice with water and 100 mM iron(III) chloride was applied three times. The iron-NTA resin was washed twice with water and once with 0.5% formic acid. Iron-NTA beads were resuspended in water to create a 25% resin slurry. 50 μl of Fe-NTA resin slurry was transferred to individual Silica C18 MicroSpin columns (The Nest Group) pre-equilibrated with 100 μl of 80% CAN, 0.1% TFA on a vacuum manifold. Subsequent steps were performed with the Fe-NTA resin loaded on the Silica C18 columns.

Peptide samples were mixed twice with the Fe-NTA resin and allowed to incubate for 2 minutes. The resin was rinsed four times with 200 μl of 80% ACN, 0.1% TFA. In order to equilibrate the chromatography columns, 200 μl of 0.5% formic acid was applied twice to the resin and columns. Peptides were eluted from the resin onto the C18 column by application of 200 μl of 500 mM potassium phosphate, pH 7.0. Peptides were washed twice with 200 μl of 0.5% formic acid. The C18 columns were removed from the vacuum manifold and eluted twice by centrifugation at *1000g* with 75 μl of 50% ACN, 0.1% TFA. Peptides were dried with a centrifugal adaptor and stored at −20°C until analysis by liquid chromatograph and mass spectrometry.

### Di-glycine peptide enrichment by immunoprecipitation

Peptide samples were subjected to ubiquitin remnant immunoaffinity purification with 31 μg of ubiquitin remnant antibody (Cell Signaling). Peptides were lyophilized for two days to remove TFA in the elution. The lyophilized peptides were resuspended in 1 ml of IAP buffer (50 mM 4-morpholinepropnesulfonic acid, 10 mM disodium hydrogen phosphate, 50 mM sodium chloride, pH 7.2). Peptides were sonicated and centrifuged for 5 minutes at 16,100g. Ubiquitin remnant beads were washed twice with IAP buffer and incubated with the peptides at 4°C for 90 minutes with rotation. Unbound peptides were separated from the beads after centrifugation at 700g for 60 seconds. Beads containing peptides with di-glycine remnants were washed twice with 500 μl of water and peptides were eluted twice with 55 μl of 0.15% TFA. Di-glycine remnant peptides were desalted with UltraMicroSpin C18 column (The Nest Group). Desalted peptides were dried with a centrifugal adaptor and stored at −20°C until analysis by liquid chromatograph and mass spectrometry.

### Liquid chromatography and mass spectrometry

Peptides were analyzed in technical duplicate on a ThermoFisher Orbitrap Fusion Lumos Tribid mass spectrometry system equipped with an Easy nLC 1200 ultrahigh-pressure liquid chromatography system interfaced via a Nanospray Flex nanoelectrospray source. Samples were injected on a C18 reverse phase column (25 cm x 75 μm packed with ReprosilPur C18 AQ 1.9 um particles). Peptides were separated by an organic gradient from 5 to 30% ACN in 0.02% heptafluorobutyric acid over 180 min at a flow rate of 300 nl/min for the phosphorylated peptides or unmodified peptides for global abundance. Ubiquitinated peptides were separated by a shorter gradient of 140 min. Spectra were continuously acquired in a data-dependent manner throughout the gradient, acquiring a full scan in the Orbitrap (at 120,000 resolution with an AGC target of 400,000 and a maximum injection time of 50 ms) followed by as many MS/MS scans as could be acquired on the most abundant ions in 3 s in the dual linear ion trap (rapid scan type with an intensity threshold of 5000, HCD collision energy of 32%, AGC target of 10,000, maximum injection time of 30 ms, and isolation width of 0.7 *m/z*). Singly and unassigned charge states were rejected. Dynamic exclusion was enabled with a repeat count of 2, an exclusion duration of 20 s, and an exclusion mass width of ±10 ppm.

### Label-free quantitation and statistical analysis

Mass spectrometry data was assigned to mouse sequences and MS1 intensities were extracted with MaxQuant (version 1.6.0.16)^50^. Data were searched against the SwissProt mouse protein database (downloaded on March 3, 2018). Trypsin (KR|P) was selected allowing for up to two missed cleavages. Variable modification was allowed for N-terminal protein acetylation and methionine oxidation in addition to phosphorylation of serine, threonine, and tyrosine, or diGly modification of lysine, as appropriate. A static modification was assigned to carbamidomethyl cysteine. The other MaxQuant settings were left at the default.

Statistical analysis of MaxQuant-analyzed data was performed with artMS Bioconductor package (version 0.9) which performs the relative quantification using the MSstats Bioconductor package (version 3.14.1)^51^. Contaminants and decoy hits were removed. The samples were normalized across fractions by median-centering the log_2_-transformed MS1 intensity distributions. The MSstats group comparison function was run with no interaction terms for missing values, no interference, unequal intensity feature variance, restricted technical and biological scope of replication. Log_2_(fold change) for protein/sites with missing values in one condition but found in > 2 biological replicates of the other condition of any given comparison were estimated by imputing intensity values from the lowest observed MS1-intensity across samples^25^; *p* values were randomly assigned between 0.05 and 0.01 for illustration purposes. Statistically significant changes were selected by applying a log_2_-fold-change (>1.0 or <-1.0) and an adjusted *p* value (<0.05) corrected for multiple comparison.

Phosphopeptide data was analyzed with MSstats using the unmodified output data from MaxQuant and phosphosite collapsed data. Phosphopeptide data was collapsed using an in-house script that assigns phosphopeptide sites to the most likely phosphorylated residue in peptides where multiple phosphorylated residues were detected with low phosphorylation probability scores.

### Gene enrichment analysis

Gene enrichment analysis was performed with Metascape (metascape.org) with Custom analysis settings using only GO Biological Processes. Heatmaps were made with the R-based Complex heat map program and the Ward D clustering algorithm.

### PhosFate analysis

Murine phosphorylation sites were mapped to the human proteome using R. Phosphopeptide log_2_-fold-change profiles were uploaded to the PhosFate Profiler tool (Phosfate.com^33^) for kinase and kinase complex predictions.

### RNAseq

Bone marrow-derived macrophages were infected at a MOI of 1 for the 24-hour time point and a MOI of 10 for 2- and 6-hour time points. At the indicated time points, monolayers were washed with PBS, and macrophages were resuspended in 1 ml of TRIzol (ThermoFisher). Following the addition of chloroform (200 μl), samples were mixed and centrifuged for 10 min. at 4 °C. An equal volume of 70% ethanol was added to the aqueous layer, and RNA was extracted using silica spin columns (Invitrogen PureLink RNA Mini Kit). Purified RNA was treated DNAse (New England Biolabs) followed by EDTA. RNAseq libraries were prepared from biological triplicate samples by the University of California, Davis, Expression Analysis Core. Differential gene expression analysis was performed by the University of California, Davis, Bioinformatics Core. Transcriptional changes of similar levels between the 24-hour datasets (standard deviation <1 for the log_2_fold changes) measured in our laboratory and a previously published study of *Mtb* infected bone derived macrophages were included in the analysis.

### Cloning and lentiviral transduction of FLAG-tagged autophagy receptors

PCR products were generated for each autophagy adaptor using the following primer pairs: P30/P2 (p62), P31/P4 (BNIP3), P32/P6 (BNIP3L), P33/P8 (FUNDC1). The remaining autophagy receptors were cloned by splicing with overlap extension PCR with the following primers: P34/P37 and P38/P10 (NBR1), P35/P39 and P40/P12 (Optineurin), P36/P41 and P42/P14 (TAX1BP1). PCR products were cloned into an entry vector encoding a N-terminal triple FLAG tag (pENTR-N-FLAG; Parry *et al.,* in preparation) using sequence- and ligation-independent cloning. The Gateway LR reaction was used to introduce the cloned autophagy receptors into the pLenti CMV Puro DEST destination vector^52^. RAW264.7 cells were transduced and subjected to puromycin selection for 7 days.

### Cloning and lentiviral transduction of CRISPR guide RNAs

Single oligonucleotide guides for three scramble controls and two exons in *Tax1bp1* selected from the Brie library were closed into lentiGuide puro (Addgene #52963) using the following primer pairs: P174/P175 (Scramble 1), P176/P177 (Scramble 2), P178/P179 (Scramble 3), P160/P161 (*Tax1bp1* exon 10), and P164/P165 (*Tax1bp1* exon 9). 293T cells were reverse transfected with the vectors encoding scramble or *Tax1bp1* guides. Bone marrow cells from Rosa26-Cas9 knock-in mice were transduced with retrovirus and differentiated in the presence of puromycin. Transduced and fully-differentiated bone marrow-derived macrophages were cultured for three days in the absence of puromycin before seeding for microscopy experiments.

### Western blot

The anti-FLAG M2 mouse monoclonal antibody was purchased from Sigma-Aldrich. 60 μg of protein lysate was separated by SDS-PAGE (BioRad Miniprotean TGX 4-20%) and transferred onto nitrocellulose membranes. After probing with the anti-FLAG M2 antibody at a dilution of 1:1000, membranes were imaged on an Odyssey scanner (Li-cor).

In order to probe for actin on the same nitrocellulose membrane, antibodies were stripped from the membrane with 0.2 N sodium hydroxide and re-probed with the β-actin mouse monoclonal antibody (C4; Santa Cruz Biotechnology, SC-47778).

### Immunofluorescence microscopy for FLAG-tagged autophagy receptor colocalization

Infected macrophages were washed with PBS and incubated with blocking and permeabilization buffer (0.05% saponin, 5% fetal calf serum, PBS) for 30 minutes at room temperature. Coverslips were incubated with anti-FLAG M2 mouse monoclonal antibody at a dilution of 1:500 for 3 hours at room temperature and secondary antibody conjugated to fluorophore at a dilution of 1:1000 (Invitrogen). 45 images per coverslip were obtained with a Nikon Eclipse TE2000-E pinning disc confocal microscope equipped with an Andor laser system and Borealis beam conditioning unit. Images were obtained with the 100X oil objective for figures and 40X air objective for colocalization.

### Immunofluorescence microscopy for p62, ubiquitin, and phospho-TBK1

Immunostaining for p62 and ubiquitin was performed with blocking and permeabilization buffer (1% saponin, 3% bovine serum albumin) and anti-p62 rabbit monoclonal antibodies (Abcam #AB109012) at a concentration of 1 μg/ml or anti-ubiquitylated protein antibodies (clone FK2, Millipore Sigma #04-263) at a dilution of 1:400 for 3 hours at room temperature.

Immunostaining for phospho-TBK1 was performed with blocking and permeabilization buffer (0.3% Triton X-100, 1% bovine serum albumin in PBS) and anti-phospho-Ser172-TBK1 (D52C2) XP Rabbit monoclonal antibody (Cell Signaling #5483) for three hours at room temperature. Samples were incubated with a secondary antibody conjugated to Alexa Fluor 488 fluorophore at a dilution of 1:4000 (Invitrogen) and DAPI at a dilution of 1:1000 for one hour at room temperature. 25 images per well were obtained with a Perkin Elmer Opera Phenix High Content Screening confocal microscope at 40X magnification for colocalization or 63X for figures. In the images shown from the Rosa26-Cas9 knock-in bone marrow-derived macrophages, low-level background GFP fluorescence was filtered out. Automated colocalization measurements were performed with the Perkin Elmer Harmony software package. A pipeline was created to measure colocalization of *Mtb* and autophagy markers (p62, FK2, and phospho-TBK1) in macrophages infected with two *Mtb* bacilli.

### Immunofluorescence microscopy for LC3

Immunostaining for LC3 was performed with blocking and permeabilization buffer (0.3% Triton X-100, 2% bovine serum albumin) and anti-LC3 (clone 2G6, Nanotools #0260-100/LC3-2G6) at a dilution of 1:200 for 3 hours at room temperature. Samples were incubated with a secondary antibody conjugated to Alexa Fluor 647 fluorophore at a dilution of 1:1000 (ThermoFisher) and DAPI at a dilution of 1: 1000 for one hour at room temperature. 39 images per well were obtained with the Opera Phenix confocal microscope at 40X magnification for colocalization or 63X for figures. Automated colocalization measurements were performed in macrophages infected with two *Mtb* bacilli.

## Acknowledgements

We acknowledge members of the Cox and Krogan labs for their comments on this manuscript submission. We thank H. Skip Virgin and Hidekatsu Iha for *p62^−/−^* and *Tax1bp1^−/−^* mice, respectively. We thank Lori Kohlstaedt at the UC Berkeley proteomics core for assistance in LC-MS analysis of our peptide samples, and Christopher Noel for assistance in the development of the Perkin Elmer Harmony microscopy colocalization pipelines. This work was supported by NIH grants P01 AI063302 (N.J.K, J.S.C.), P50 GM082250 (N.J.K), U19 AI106754 (N.J.K), DP1 AI124619 (J.S.C), and R01 AI120694 (N.J.K and J.S.C). J.M.B was supported by T32 training grant (4T32HL007185-39 & −40, Dean Sheppard PI) and NIH K12 (5K12HL119997-05, David Erle PI). Research reported in this publication was supported by the Office of The Director, National Institutes of Health of the National Institutes of Health under Award Number S10OD021828 for the Opera Phenix and associated staff support. The content is solely the responsibility of the authors and does not necessarily represent the official views of the National Institutes of Health.

---

**Fig. S1.**
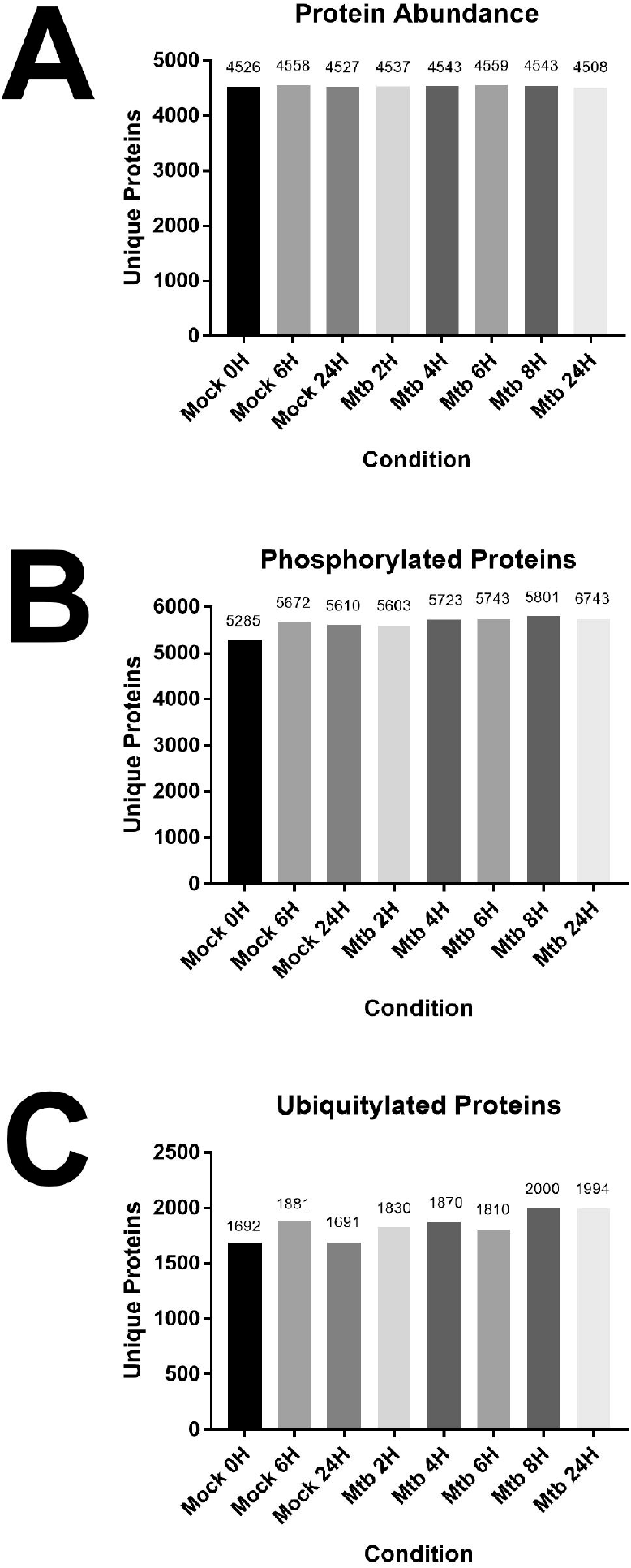
Unique host proteins identified at each time point 2-24-hours post-infection with *Mtb* or mock infection. (*A-C*) Graph of the number of unique proteins identified by mass spectrometry in biological triplicate, technical duplicate samples for each experimental condition from unenriched (*A*), phosphoenriched (*B*), or di-Gly enriched (*C*) peptide samples.

**Fig. S2.**
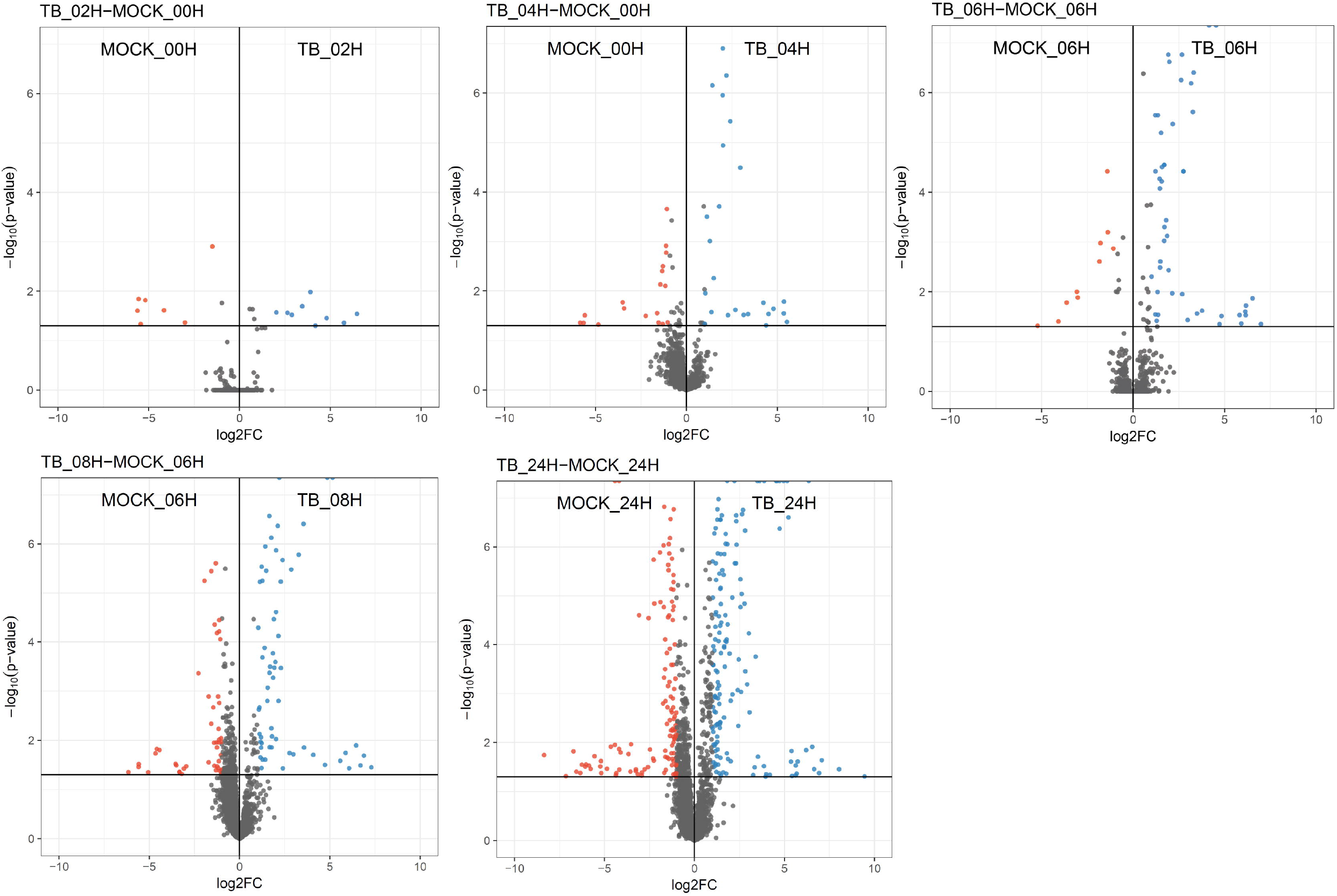
Host protein abundance changes during a time course of *Mtb* infection. Volcano plots displaying the proteins changing in abundance during *Mtb* infection. Proteins with a *p* value less than 0.05 and log_2_(fold change) greater than 1 are colored red. Proteins with a *p* value less than 0. 05 and log_2_(fold change) less than −1 are colored blue.

**Fig. S3.**
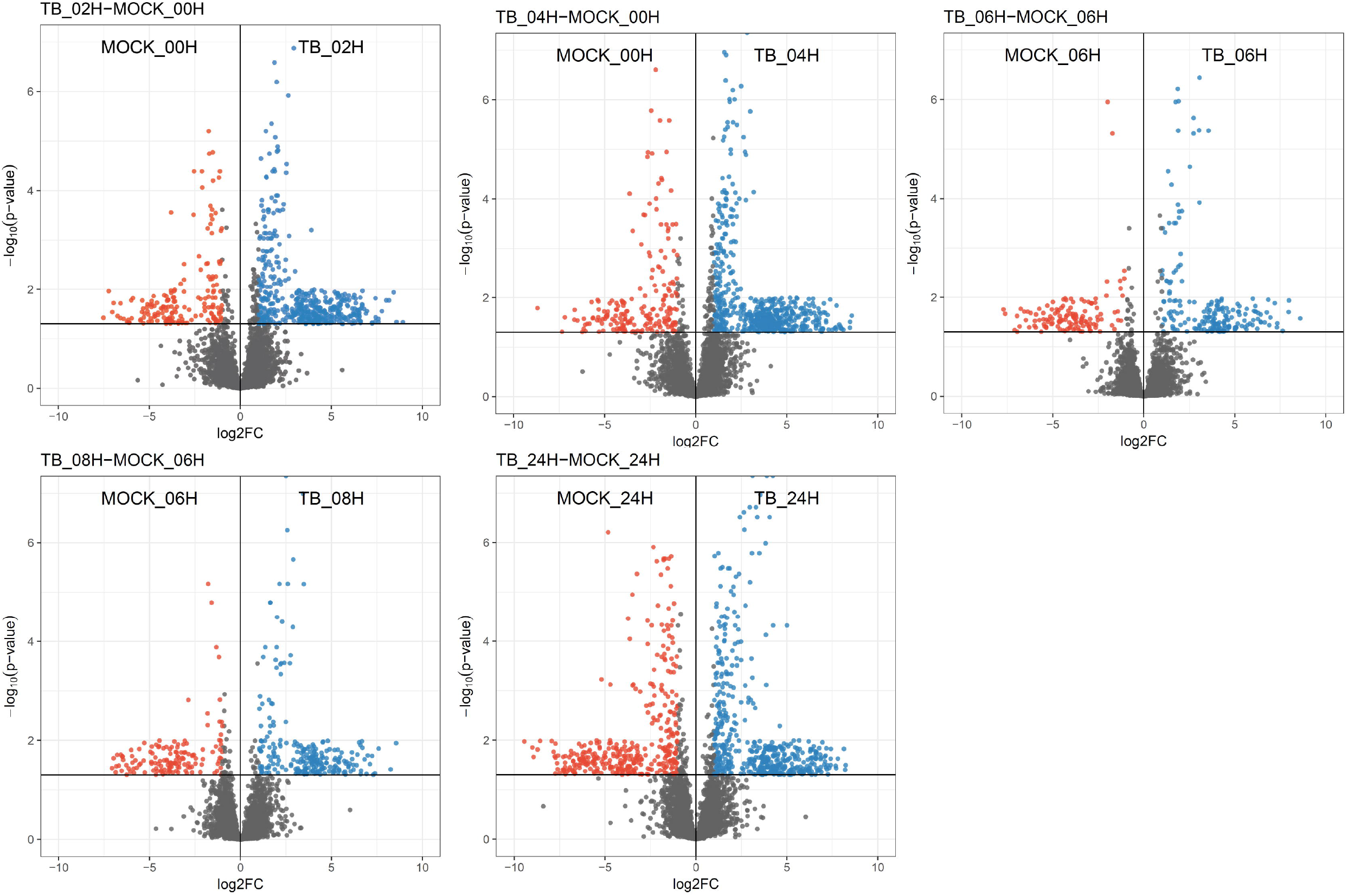
Changes in host protein phosphorylation during a time course of *Mtb* infection. Volcano plots displaying the changes in phosphopeptides during *Mtb* infection. Phosphopeptides with a *p* value less than 0.05 and log_2_(fold change) greater than 1 are colored red. Phosphopeptides with a *p* value less than 0.05 and log_2_(fold change) less than −1 are colored blue.

**Fig. S4.**
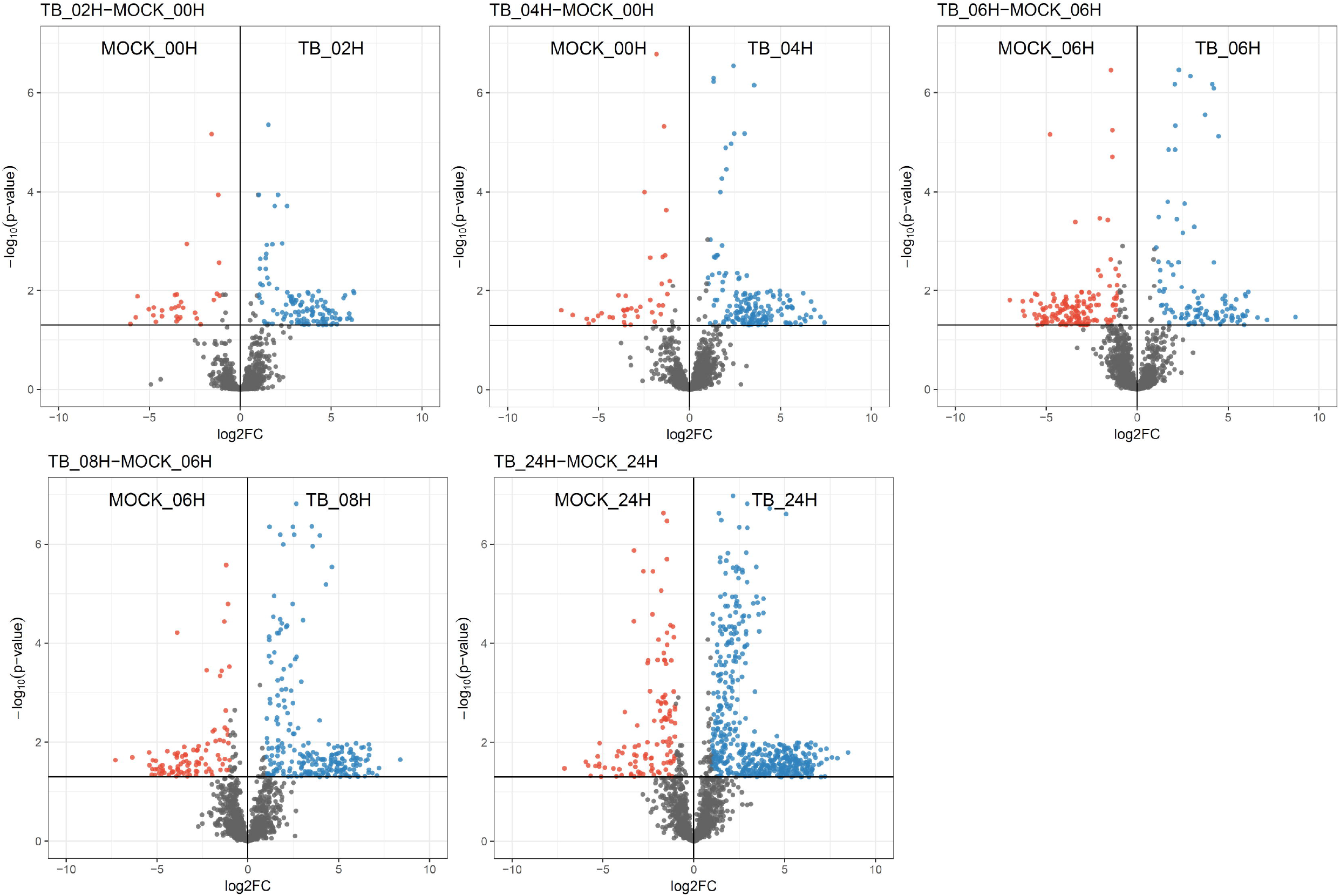
Changes in host protein ubiquitylation during a time course of *Mtb* infection. Volcano plots displaying the changes in di-Gly modified peptides during *Mtb* infection. Ubiquitylated peptides with a *p* value less than 0.05 and log_2_(fold change) greater than 1 are colored red. Ubiquitylated peptides with a *p* value less than 0.05 and log_2_(fold change) less than −1 are colored blue.

**Fig. S5.**
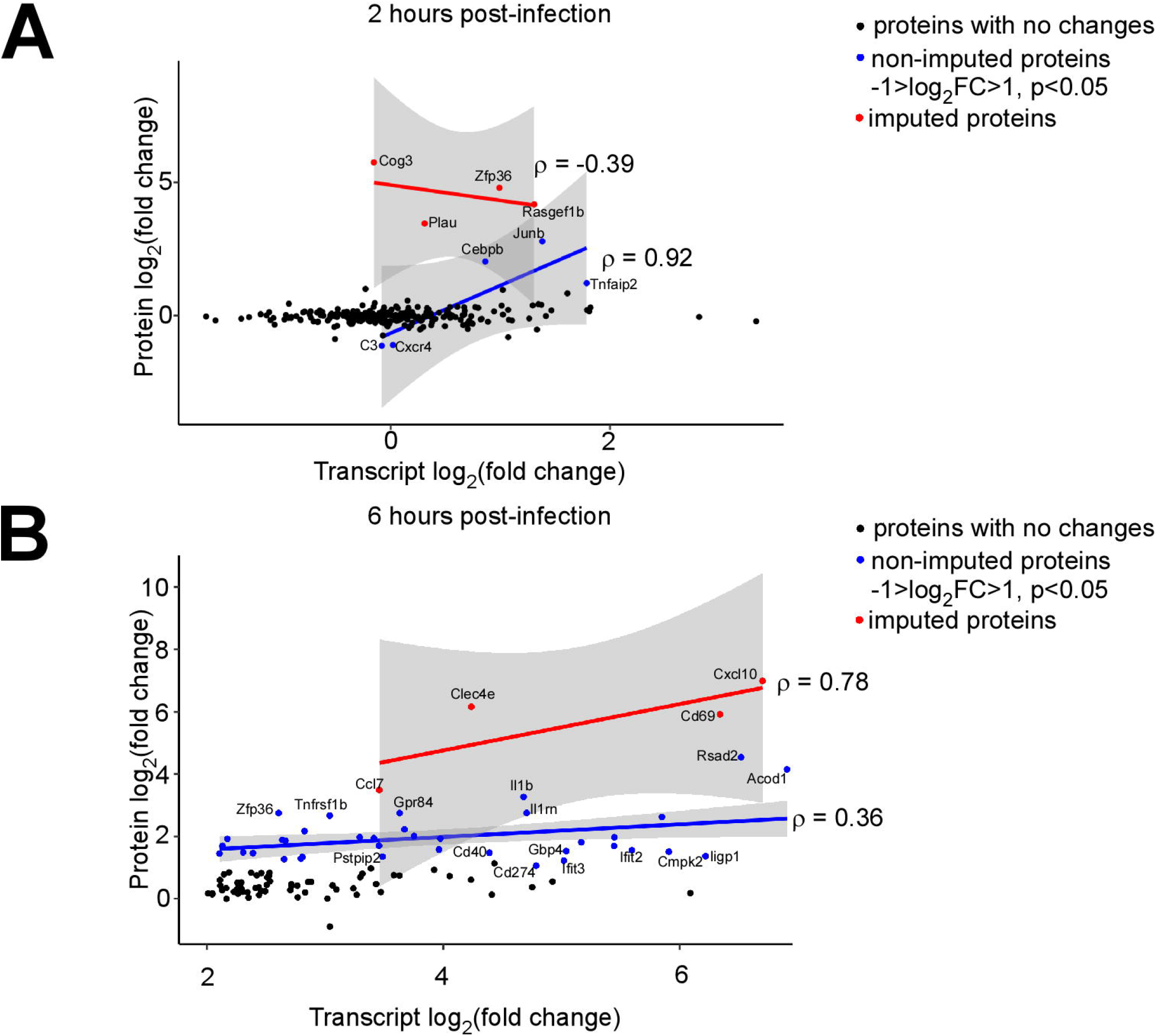
Correlation between host transcript and protein abundance changes during *Mtb* infection. (*A, B*) Correlation plot of changes in gene transcription and protein abundance 2- (*A*) and 6-hours (*B*) post-infection with *Mtb*. Proteins with no statistically significant changes during *Mtb* infection are colored black. Non-imputed proteins with statistically significant changes during *Mtb* infection are colored blue. Imputed proteins during *Mtb* infection are colored red. The correlation coefficient for non-imputed and imputed proteins are both shown.

**Fig. S6.**
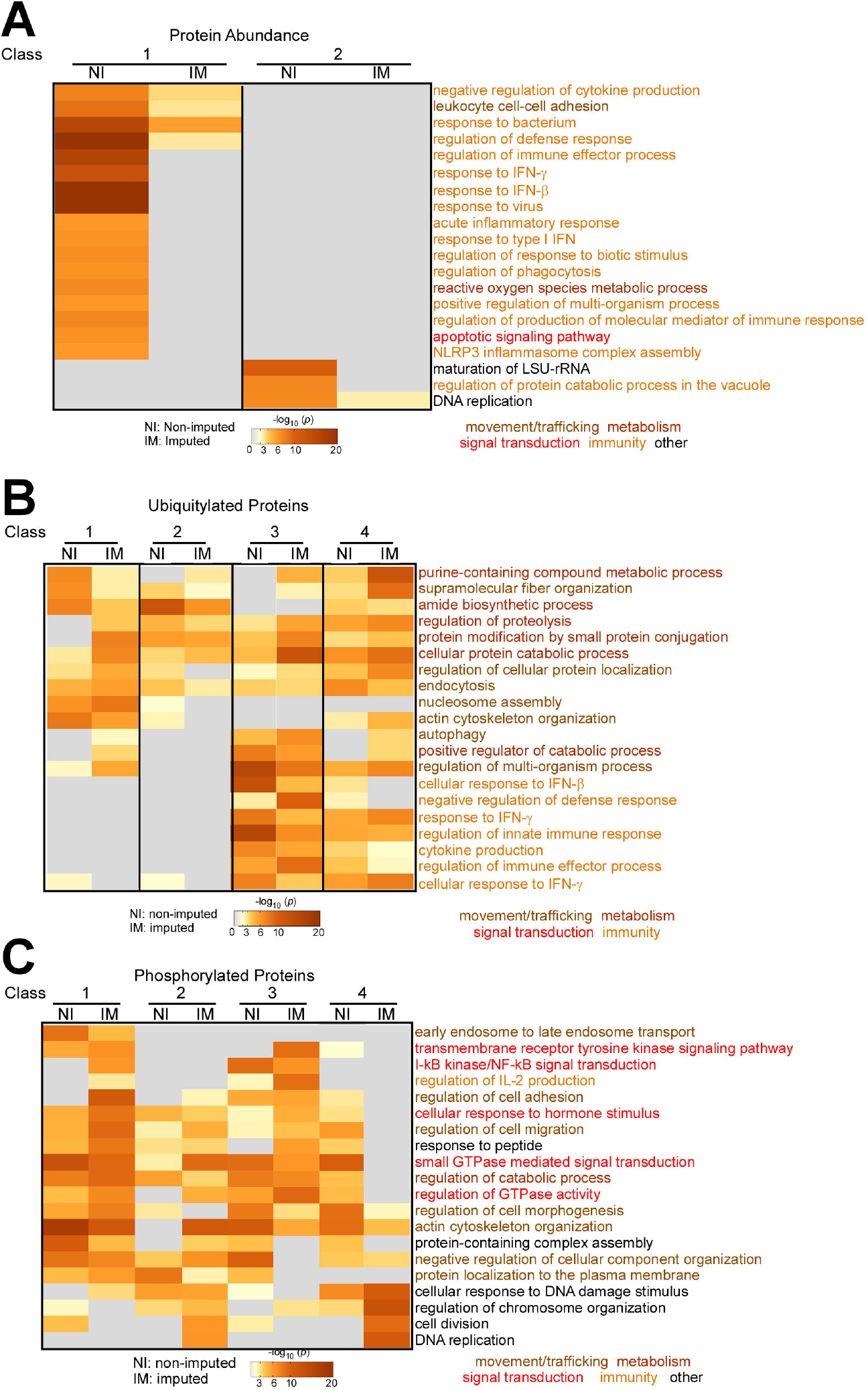
Functional enrichment terms of host proteins changes during *Mtb* infection. (*A-C*) Heat map displaying functional enrichment terms for the classes of protein changes (Fig. 3-5) during *Mtb* infection. Non-imputed (NI) and imputed (IMP) measurements were separated for the analysis.

**Table S1.** R values for peptide intensity levels from biological and technical replicates.

**Table S2.** Primers used in this study.

**Table S3.** Autophagy receptors changing in protein abundance, phosphorylation, or ubiquitylation during *Mtb* infection.

**Table S4.** Protein abundance changes during *Mtb* infection.

**Table S5.** Changes in phosphorylation during *Mtb* infection.

**Table S6.** Changes in ubiquitylation during *Mtb* infection.

